# On the involvement of cell volume regulation mechanisms in post-hypertonic lysis and slow-freezing injury of human erythrocytes and its broader cryobiological significance

**DOI:** 10.1101/2023.04.28.538673

**Authors:** Ivan Klbik

## Abstract

An effort is made to explain the salt loading of hypertonicity-treated human red blood cells (RBC) and their post-hypertonic lysis, which occurs upon resuspension to isotonic media. It is recognized that the salt loading is consistent with the regulatory volume increase response - a physiological response applied by cells to adapt to hypertonic conditions. It is therefore hypothesized that salt loading occurs via physiological ion transport pathways, allowing passive dissipative fluxes across the plasma membrane with the preservation of membrane integrity. It is argued that post-hypertonic lysis is essentially a hypotonic lysis occurring already in an isotonic medium due to the excess salt in the cytoplasm. A simplified physiological model accounting for transmembrane ion fluxes was applied to simulate the RBC physiological state to address the problem quantitatively. Instead of assuming the plasma membrane to be impermeable to ions, the dynamic nature of cell physiological state and capacity of cells to perform acute cell volume regulation by increasing passive cation plasma membrane permeability is recognized. It is shown that the RBC plasma membrane contains ion pathways capable of supporting a several-order increase in catio permeability, facilitating overall salt loading. The obtained picture, consistent with general physiology, predicts the experimentally observed RBC behavior. Although treated in theory, evidence supporting the proposed theory is provided, and the high explanatory power of the proposed idea is demonstrated. While focused on human RBCs, this study offers a general mechanism for the osmotic injury of cells exposed to hypertonic solutions - conditions relevant to cell cryopreservation - since many cells possess cell volume regulation mechanisms. Implications of the proposed hypothesis explaining the cryoprotective mechanism of common cryoprotectants are discussed, and additives targeting ion transport pathways are suggested as novel cryoprotectants.

## INTRODUCTION

Human red blood cells are subject to post-hypertonic lysis, which occurs during resuspension to isotonic solution following previous exposure to hypertonic electrolyte and non-electrolyte media. The post-hypertonic lysis was first described by Takei in 1921,^1^ and later by Lovelock,^2^ Zade-Oppen,^3, 4^ and more recently by others.^5^ Post-hypertonic lysis of other mammalian RBCs was also observed.^6^ Lovelock demonstrated the post-hypertonic lysis as a dominant injury associated with exposure of RBCs to concentrated salt solutions. A correlation between the damaging salt concentration (and osmolality) attained during osmotic and freeze-thaw experiments was observed, implying the same mechanism of injury, emphasizing the cryobiological relevance of the post-hypertonic lysis. Since then, several authors have recognized post-hypertonic lysis as an aspect of RBC slow-freezing injury when cryopreserved cells are exposed to freeze-concentrated salts as ice precipitates out of the solution.^7, 8^ The extent of RBCs post-hypertonic lysis was found to be proportional to the level of hypertonicity, the duration of exposure to the hypertonic media, and inversely proportional to the tonicity of the resuspension medium.^3^ Later, post-hypertonic lysis of other human cells – granulocytes,^9^ spermatozoa,^10^ and prostatic adenocarcinoma cells (PC-3)^11^ was also observed. Söderström^12^ and Lovelock^2^ proposed that the RBC plasma membrane could become permeable to sodium ions under hypertonic conditions. They argued that increased cytoplasmic sodium concentration would lead to osmotic water intake, resulting in cell swelling and lysis during the resuspension to isotonic media if a critical salt concentration and associated cell volume were exceeded. The RBC plasma membrane was indeed found to be permeable to ions under hypertonic conditions when sodium influx and potassium outflux were observed,^13–19^ supporting the previous speculations. An inward sodium leak is already occurring at 1000 mOsm/kg without simultaneous hemolysis,^15^ while an outward potassium leak occurs only above 1800 mOsm/kg - correlating with the onset of post-hypertonic lysis.^14, 17^ To explain the salt loading, Lovelock suggested that the increased ion permeability might be caused by plasma membrane changes induced by increased ionic strength. Meryman,^16, 17^ hypothesized that RBCs could shrink only to a specific limit, with further increase of tonicity resulting in the development of an osmotic pressure gradient, which is diminished by intake of extracellular osmotically active solutes. Meryman suggested that the ion leaks occur by reversible loss of plasma membrane integrity and that post-hypertonic lysis results from osmotic swelling, as previously hypothesized by Lovelock and Söderström. However, Meryman’s hypothesis on the development of osmotic pressure gradient was disproven when RBCs were shown to remain in osmotic equilibrium under the studied hypertonic conditions.^14^

In the past, when most of the cited studies about post-hypertonic lysis of RBCs were conducted, an accurate physiological picture was still developing. The authors of the mentioned studies assumed the RBC plasma membrane to be normally impermeable to cations. Due to a lack of knowledge of cell physiology and membrane ion transport during the publication of these studies, the authors did not recognize the possible involvement of membrane ion transport in their accounts, so they could not provide a physiological route for salt loading. Only Muldrew, in 2008, proposed that sodium flows into the cells via ion channels to replace the potassium that binds to salted-in proteins due to increased salt concentrations.^20^ Cell physiology now recognizes the ability of biological cells to induce cation permeability alterations, which could facilitate salt loading of human RBCs. Such a response would be consistent with cells’ regulatory volume increase (RVI) actions to adapt to hypertonic conditions. However, there is no account of post-hypertonic lysis considering the physiological perspective with an explicit account of transmembrane ion fluxes, including consideration of plasma membrane electric potential. In cryobiological descriptions, it is often assumed that the cell plasma membranes are impermeable to ions. Even beyond RBCs, only a few studies have considered the potential role of cell volume regulation in cryobiologically relevant situations. That includes the work of Peckys and Mazur,^21^ who emphasized the possible involvement of CVR at certain stages of cell handling (especially before cryopreservation during the loading of cryoprotectants) but doubted the involvement during actual cryopreservation due to an inhibitory effect of reduced temperature upon ion transport rates. However, they considered only the participation of active ion transport and did not recognize the possible role of passive dissipative ion fluxes, which have a decisive role in acute cell volume regulation.^22, 23^ Another study showed that ion transport pathways facilitating RVI response play a role in the rehydration step following cryopreservation of HepG2 and HeLa cells and that activators of these pathways affect resulting cell viability,^24^ supporting the involvement of cell volume regulation during cryopreservation, emphasizing the importance of physiological perspective in cryobiology.^25^

Based on these considerations, this work examines the possible role of cell volume regulation mechanisms in salt loading and post-hypertonic lysis of human RBCs. Based on the established physiological knowledge, it is hypothesized that salt loading is caused by an RVI response mediated by ion-specific membrane pathways involved in cell volume regulation – without any assumption of the formation of membrane lesions under hypertonic conditions or during resuspension to isotonic media. It is also hypothesized that post-hypertonic lysis is a direct cause of the increased intracellular salt content leading to the osmotic equilibration at a cell volume higher than normal isotonic RBC volume. Therefore, it is argued that post-hypertonic lysis is essentially a hypotonic lysis occurring at a lower-than-normal osmolality of the bathing solution due to the accumulated salt (see Figure 1). This study aims to evaluate this hypothesis by applying a quantitative model based on the pump-leak concept,^26^ which, instead of assuming the RBC plasma membrane to be impermeable to cations, recognizes the dynamic equilibrium of normal physiological cell steady state, explicitly addressing the transmembrane ion fluxes with account for plasma membrane electric potential.

**Figure 1.**
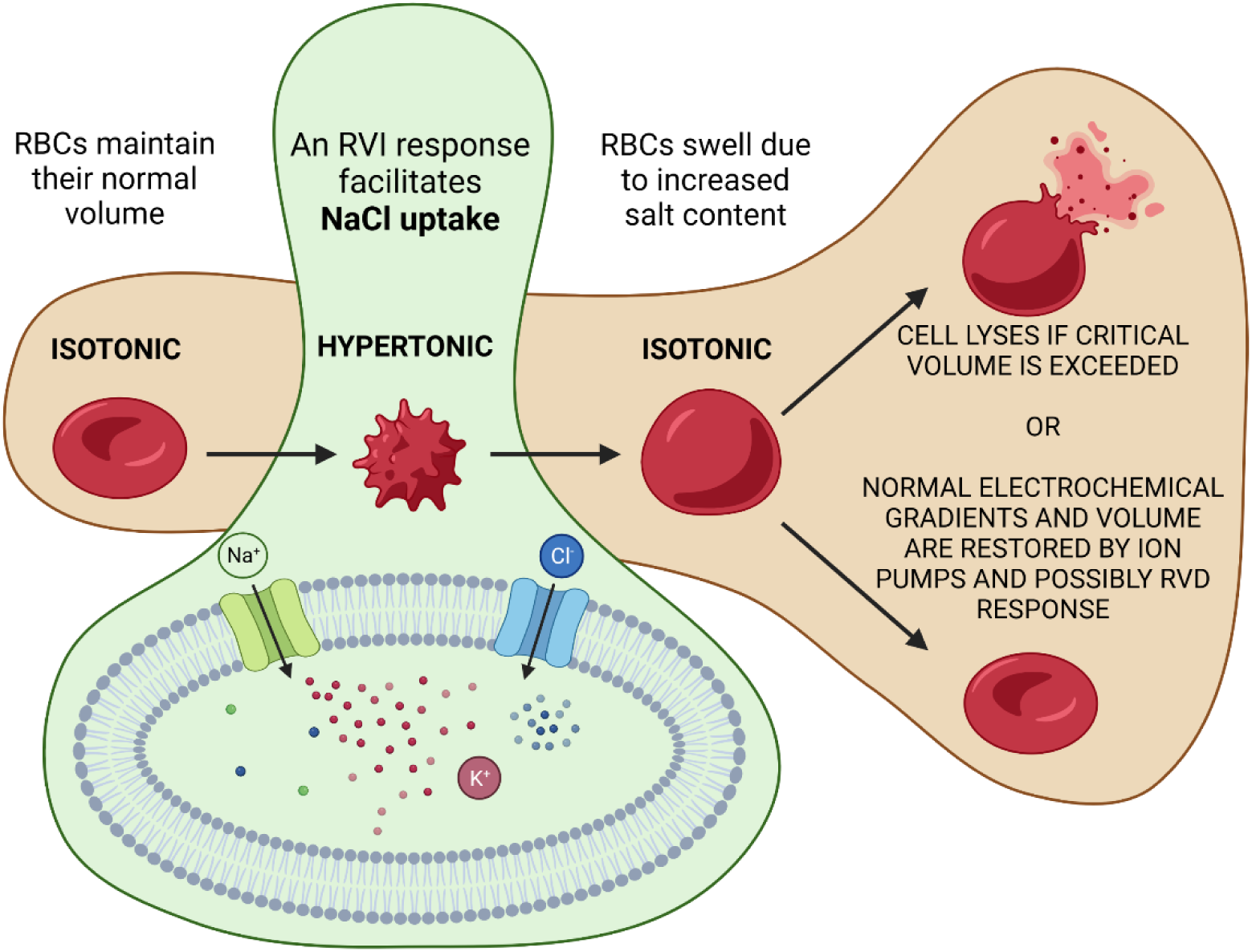
A picture illustrating the proposed hypothesis, which claims that salt loading of hypertonicity-treated human RBCs is caused by a regulatory volume increase response initiated by RBCs to adapt to anisotonic conditions. Although this response is a healthy physiological reaction to changing conditions, the detrimental aspect of this response lies in an excessive salt accumulation that causes osmotic lysis of RBC upon abrupt resuspension in an isotonic medium. A more detailed picture, including specific ion transport pathways, is shown in Figure 7.

## CELL PHYSIOLOGY AND CELL VOLUME REGULATION

To fully appreciate the hypothesis proposed in this work, a general physiological picture must be provided first to understand the normal physiological state of biological cells and possible cell response to anisotonic conditions (for introduction to cell volume regulation, readers are advised to consult review by Lang,^22^ Hoffman et al.,^27^ and first chapter in a book by Hallows and Knauf^28^). The assumption of plasma membrane impermeability to ions has often been introduced into the models describing the osmotic response of various cells. The models are developing, as evidenced by recent efforts to account for deviations in the osmotic behavior of different cells ^29–31^ by incorporating the possibility of ion movements into the mathematical models describing the cell behavior.^32–34^ It should be emphasized that although the impermeability assumption is often adequate, it is inconsistent with the current state of physiological knowledge, which perceives the plasma membrane as permeable to ions such as Na^+^, K^+^, and Cl^-^. A typical cell (including human RBC) is characterized by low intracellular content of sodium ions and high content of potassium ions and by negative electric membrane potential at steady-state conditions. Human RBCs are also characterized by relatively high permeability to passively distributed chlorine ions (about two orders higher than the normal cation permeability)^35^ and associated high intracellular chlorine content (∼75 mmol/L) with an equilibrium membrane potential of about -10 mV.^36^ While animal cells are normally in osmotic equilibrium at static physiological conditions, they are not in chemical equilibrium as a thermodynamic driving force is present for sodium ions to enter and for potassium ions to escape the cell (see Figure 2). These electrochemical gradients and cell volume are maintained by ion pumps counteracting passive dissipative ion fluxes across the plasma membrane, making the membrane effectively impermeable to these ions – emphasizing the dynamic nature of the normal physiological steady state. Therefore, the assumption of plasma membrane ion impermeability is adequate until the dynamic equilibrium between the influx and outflux of these ions holds.

**Figure 2.**
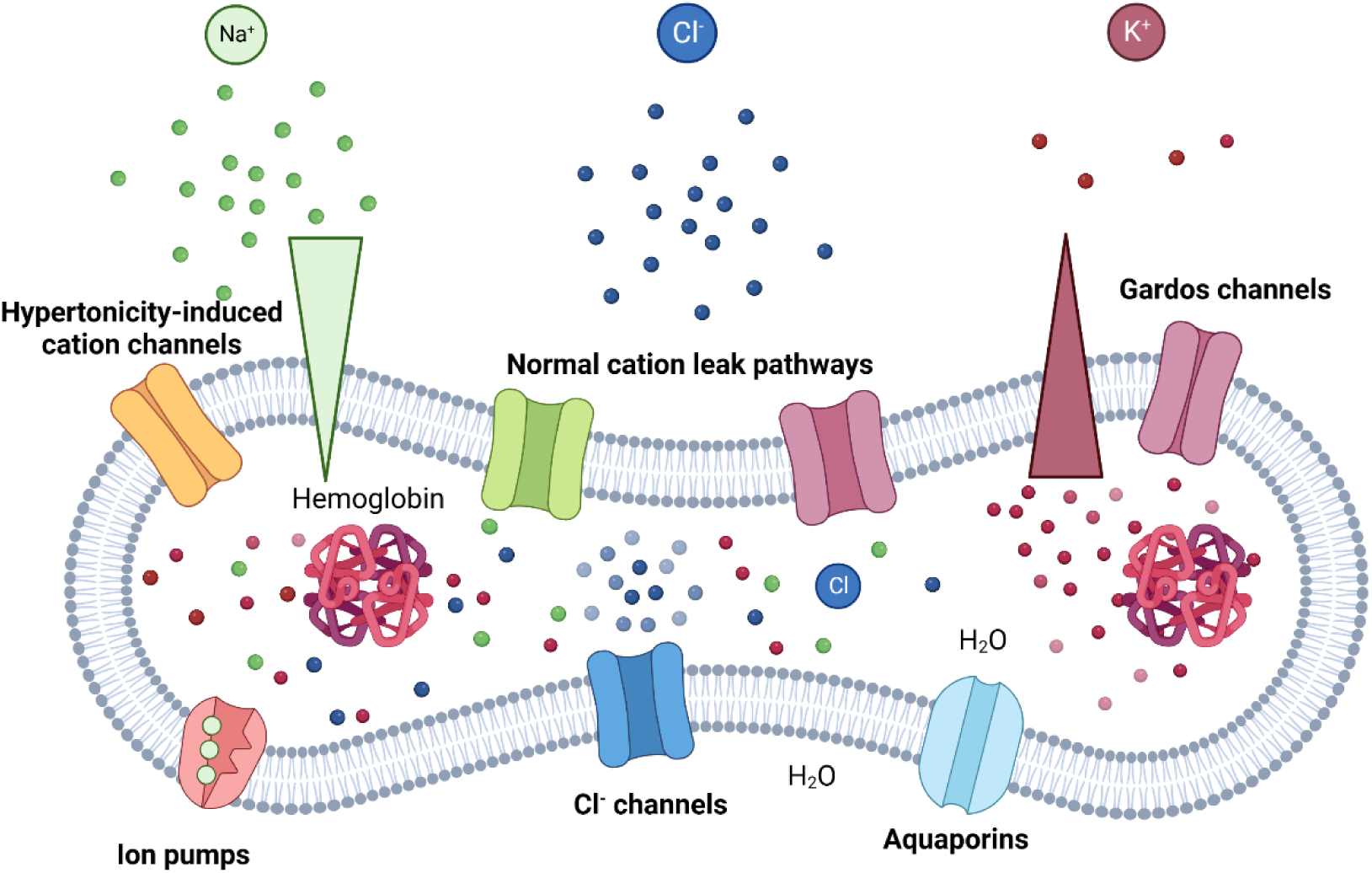
A picture illustrating the dynamic nature of cell steady-state when dissipative ion fluxes are counteracted by ion pumps (actual RBC plasma membrane contains various ion transporters and ion channels, see for example, the book by Scheer et al.37). The presence of electrochemical gradients for sodium and potassium ions is emphasized, which might by harvested to perform cell volume regulation by activation of pathways normally silent at normal isotonic conditions.

Electrochemical gradients for the pumped cations allow cells, besides performing various physiologically essential functions, to respond and adapt to changing conditions by allowing acute cell volume regulation^38^ (for the broader functional significance of cell volume regulation mechanisms, consult a review by Lang et al.^39^). Cell volume regulation mechanisms are broadly categorized into regulatory volume increase and regulatory volume decrease (RVD) actions. RVI mechanisms facilitate uptake, while RVD induces loss of osmotically active solutes into the cytoplasm, followed by water influx and outflux, respectively. Cell volume regulation mechanisms allow cells to adapt to changing conditions, reduce the stress from hypo- and hypertonicity, or prevent osmotic lysis. More recently, cell volume regulation mechanisms were shown to be involved in cell proliferation and apoptosis (a suicidal form of cell death), characterized by (isotonic) cell swelling and shrinkage.^40–42^ The relevant time scale of the onset of cell volume regulation mechanisms varies from seconds to days. It is critical to distinguish between the acute and chronic volume regulation strategies applied to restore the original volume. The chronic volume regulation strategies include, besides the action of ion pumps, mechanisms increasing and decreasing the number of organic osmolytes. Osmolyte accumulation is performed by their synthesis, increased uptake, or decreased outflux or degradation, while their (increased) degradation or induced outflux accomplishes loss of organic osmolytes.^38^ Long-term exposure to anisotonic conditions also induces changes in the expression of proteins facilitating membrane transport.^27^ While synthesis of organic osmolytes may take hours to days,^43^ the acute cell volume regulation machinery, composed of various ion transport pathways, is activated and completed more quickly, usually within tens of minutes following the introduction of the anisotonicity.^27, 44, 45^ Considering the normal ion contents of a typical biological cell, an acute RVD is facilitated by passive loss of intracellular KCl, while passive gain of extracellular NaCl facilitates an acute RVI response.

Acute cell volume regulation is facilitated by various leak pathways formed by ion channels and transporters, which allow relatively high ion flux rates down their electrochemical gradients. Normal RBC plasma membrane is enriched with a wide variety of membrane proteins. Some of them serve as ion transport pathways giving RBCs the capacity to perform acute cell volume regulation (both RVI and RVD) to alter their physiological state (readers are advised to consult works by Scheer et al.,^37^ Kaestner,^46^ or Thomas et al.^47^ to obtain a picture of ion transport mechanisms present in RBC plasma membrane). Although the normal basal RBC cation flux rates are relatively low, activation of the normally silent ion transport pathways might dramatically increase the flux rates and facilitate acute volume regulation. Ion channels generally offer higher flux rates than ion transporters, implying a higher capacity for acute cell volume regulation. Indeed, a class of hypertonicity-induced cation channels was identified as the most efficient mechanism of regulatory volume increase actions, allowing higher permeation rates than ion transporters.^48^ The presence of shrinkage-activated cation channels (classified as HICCs) in the RBC plasma membrane has been confirmed.^49^ The capacity of human RBCs to substantially increase the cation permeability was reported several times^49–52^ with even several-order increases being observed.^53^ It is suggested here that HICCs might be a major contributor to the RVI response, which is hypothesized to cause sodium influx (coupled with an influx of chlorine ions) under hypertonic conditions – emphasizing the physiological route for the salt loading. An involvement of ion transporters such as Na^+^/K^+^/2Cl^-^ cotransporter and Na^+^/H^+^ exchanger might also be expected.^49^ A quantitative description is developed to test the proposed hypothesis by evaluating the effect of increased cation permeability on the physiological state of human RBCs, described in the following section.

## MODEL

Several models have been applied to describe the physiological state of biological cells. To name a few, Goldman-Hodgkin-Katz (GHK) equations^54, 55^ and their extension by Mullins-Noda,^56^ or Armstrong^57^ should be mentioned (for their review, readers are advised to consult the work of Fraser and Huang^58^). In this study, a charge difference method was chosen, an approach that was developed more recently and is more suitable to address the dynamic evolution of the cell’s physiological states.^26^ This model is not based on the assumption of a steady state when time changes of membrane potential are allowed, in contrast with the GHK approach. The predictions of the charge difference method have been validated against experimental data.^59^ This method represents an iterative approach that explicitly evaluates transmembrane ion fluxes with associated changes in intracellular ion contents at each iteration step. The resulting net ion flux across the plasma membrane is given by passive dissipative leak and the sodium pump contribution (except chlorine ions, which are not pumped across the RBC plasma membranes). For the time change of intracellular ion concentration, the following equations were obtained:

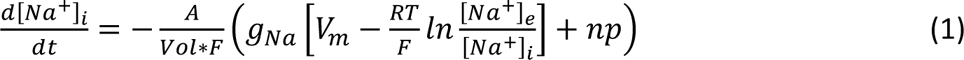

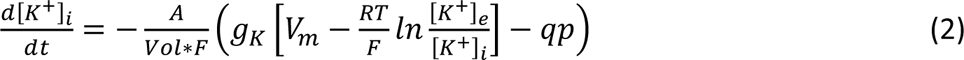

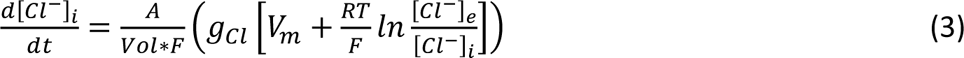

Where terms in square brackets refer to concentrations of individual ions, with subscript *i,* and *e* denoting intracellular and extracellular compartments, respectively. *g* refers to individual ion’s conductance, *A* refers to plasma membrane surface area, *Vol* refers to cell volume, *V_m_* is plasma membrane electric potential, *T* is absolute temperature, *F* and *R* are Farada’s and gas constant, respectively. *p* refers to the rate of the sodium pump, which is multiplied by *n* (equal to 3) and *q* (equal to 2) – considering the stoichiometry of the sodium and potassium ions being pumped in one cycle. The membrane potential is calculated directly from the intracellular ionic charge using the physical relationship between charge *Q*, capacitance *C*, and electric potential *V_m_*:

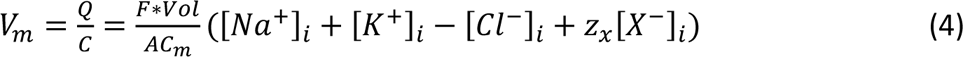

Where [*X*^−^]*_i_* refers to the concentration of intracellular non-permeants and *z_x_* refers to the valency of their charge. *C_m_* denotes the plasma membrane capacitance.

Since human RBCs are characterized by relatively high hydraulic conductivity, water movements across the plasma membrane are not addressed explicitly – an instantaneous osmotic equilibration by passive water fluxes is assumed, which causes equalization of intracellular and extracellular osmolality at all conditions:

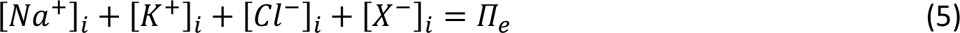

Where *π_e_* is extracellular osmolality. The following relationship then calculates the actual RBC volume:

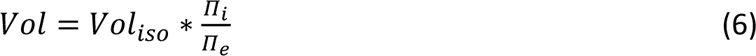

Where the ratio of intracellular and extracellular osmolality is multiplied by the normal isotonic RBC volume. The intracellular osmolality is given by intracellular concentrations assuming activity coefficients equal to one. Another implicit assumption is the infinitely diluted cell suspension when any change of intracellular ion contents does not affect the composition of extracellular fluid. This approximation is, of course, valid only at low hematocrit values of suspended RBCs. The numerical solutions of these equations were obtained by the freely available high-level programming language GNU Octave,^60^ applying the Euler method for numerical integration with a time step of one millisecond. The implementation was based on the freely available code provided in the supplementary material of a study addressing the role of ion pumps in regulating cell volume, applying the charge difference method.^61^

All parameters used in this study (Table 1 and Table 2) correspond to the human RBC state parameters at normal isotonic conditions, consistent with previous studies.^36, 50, 62, 63^ The intracellular concentration of non-permeants is assumed to be 50 mmol/L with average charge valency of -1 (indicating negative charge).^61, 64^ The only parameter that is varied in this study is the cation permeability - the quantitative consequence of the hypothesized increased permeability due to the activation of ion transport pathways facilitating acute cell volume regulation.

**Table 1.** Ion contents of RBC cytoplasm and model bathing medium.

| Ion species | Intracellular [mmol/L] | Extracellular [mmol/L] |
| --- | --- | --- |
| Na<sup>+</sup> | 22 | 151 |
| K<sup>+</sup> | 133 | 4 |
| Cl<sup>-</sup> | 105 | 155 |
| X<sup>-</sup> | 50 | 0 |

**Table 2.** Summary of the parameters used in this study.

| Parameter | Value |
| --- | --- |
| RBC surface area | 140 $\mu\text{m}^2$ |
| RBC volume | 86 fL |
| Plasma membrane capacitance | 0.1 mF/dm<sup>2</sup> |
| Sodium conductance | $2.125 \cdot 10^{-5}$ S/dm<sup>2</sup> |
| Potassium conductance | $1.416 \cdot 10^{-5}$ S/dm<sup>2</sup> |
| Chlorine conductance | $2.5 \cdot 10^{-3}$ S/dm<sup>2</sup> |
| Na/K-ATPase pump rate | $4.2135 \cdot 10^{-7}$ C/dm<sup>2</sup>s |
| Charge valency of non-permeants | -1 |

It should be noted that the normal RBC intracellular chlorine content is about 75 mmol/L in blood plasma.^36^ Note that it differs from the value assumed in this study (105 mmol/L). This study tries to mimic the conditions experienced by RBCs in the cited studies on post-hypertonic lysis, typically performed in plain sodium chloride or phosphate-buffered saline solution (low in potassium). Normal RBCs circulating in blood also contain bicarbonate and are exposed to lower extracellular Cl^-^ concentration (∼110 mmol/L) in blood plasma, resulting in an aggregate intracellular anion concentration of about 105 mmol/L. In the NaCl or buffered saline media, the intracellular Cl^-^ concentration is expected to be higher due to the absence of bicarbonate and higher extracellular Cl^-^ concentration (155 mmol/L), justifying the applied value of 105 mmol/L.

While the charge difference method is preferred for predicting the dynamic aspects of the RBC physiological state, the fundamental principle of the proposed hypothesis might be demonstrated already by consulting the steady-state approach. As it is argued, the increased cationic permeability affects the membrane potential and induces an electrochemical gradient for chlorine ions. The functional relationship between intracellular Cl^-^ concentration and steady-state cell volume was obtained as follows:^58, 61^

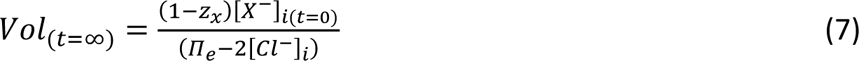

As the membrane potential determines the intracellular concentration of passive distributed chlorine ions:

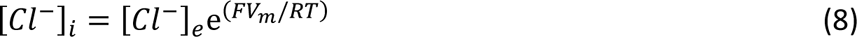

Equation (7) may be rewritten to emphasize the functional relationship between membrane potential and steady-state cell volume:

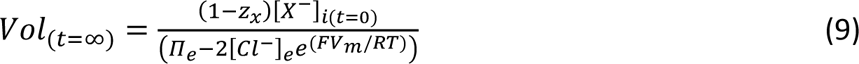

These two formulations imply that membrane depolarization and associated increased chlorine content induce isotonic cell swelling, while membrane hyperpolarization and associated decreased chlorine content have the opposite effect.

## RESULTS

The effect of increased cation conductance on membrane potential and other RBC state parameters is demonstrated in Figures 3, 4, and 5. The model predicts that increased cation conductance alters the plasma membrane potential. In the case of sodium-selective ion transport pathways, depolarization occurs - membrane potential reaches less negative values (Figure 3). On the other hand, when the potassium-selective ion transport pathways are activated, the RBC plasma membrane is hyperpolarized – a more negative membrane voltage is developed (Figure 4). In both these situations, an electrochemical gradient for chlorine ions develops. In the first case, membrane depolarization induces an inward driving force, while hyperpolarization causes an outward driving force for chlorine ions. All these simulations account for the sodium pump action, assuming the normal physiological rate. It is evident that the pump, offering only a limited capacity to extrude sodium and accumulate potassium ions, does not counteract any considerable increase in cation conductance. Therefore, increased cation conductance should dissipate normal cationic physiological electrochemical gradients. These numerical solutions explain the fundamental principle of acute cell volume regulation, as these two limiting cases (concerning cation selectivity) represent an RVI and RVD response, respectively. In the first case, an overall NaCl intake occurs, while in the second, an overall KCl loss happens, affecting the physiological state of cells – especially the cell volume. It should be noted that even in the case of cation non-selective ion transport pathways (Figure 5), a membrane depolarization eventually occurs, facilitating the entry of Cl^-^ into the cells and upregulating cell volume (consistently with previous considerations^48, 49^), although at first, some hyperpolarization and cell volume decrease is observed. An RVI associated with the activation of non-selective pathways takes more time than a purely sodium-selective variant, implying the link between the effectiveness of the cell volume regulation strategy and the selectivity of ion transport pathways triggered by hypertonicity.

**Figure 3.**
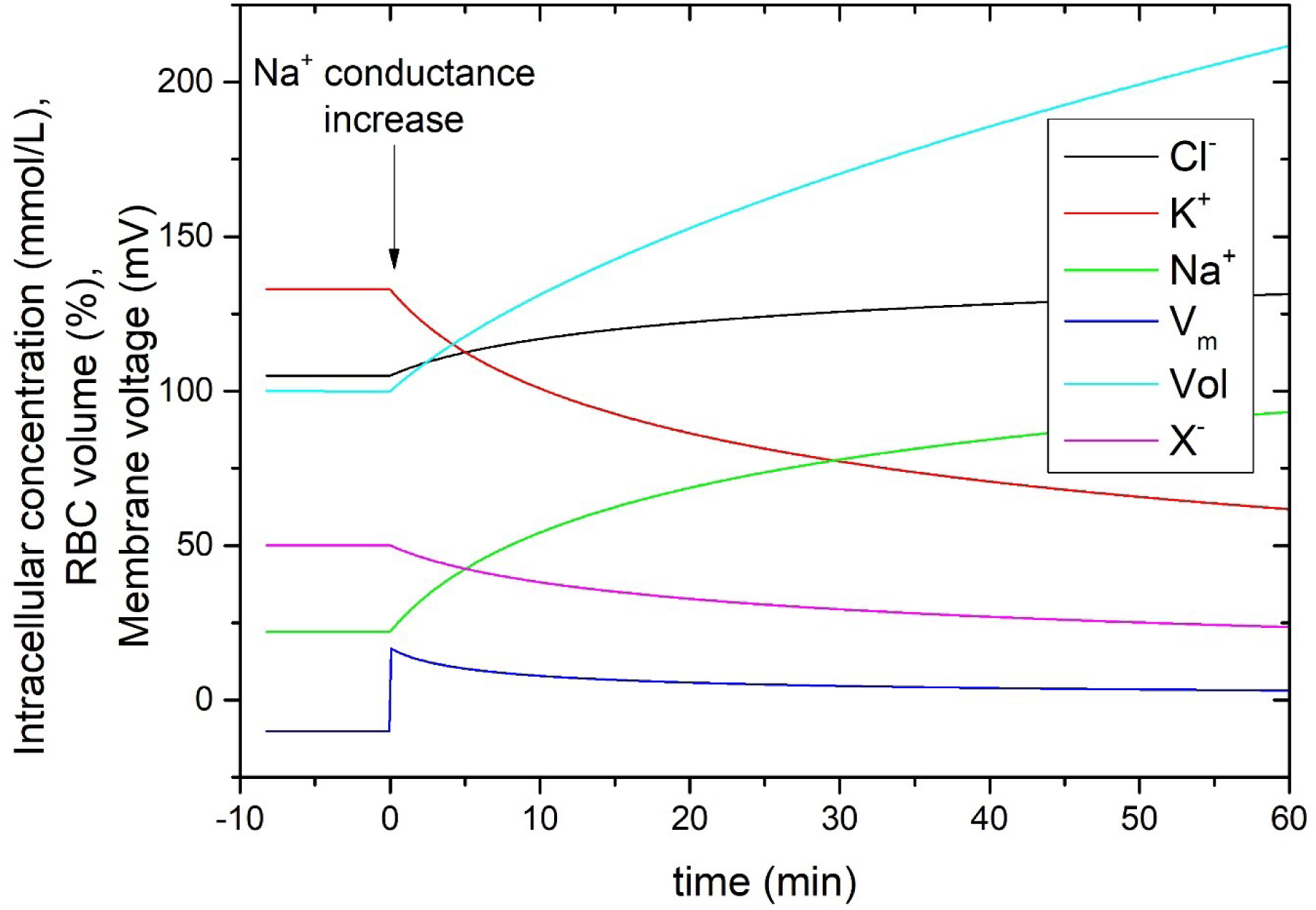
The physiological response (an RVI equivalent) to increased sodium conductance (100 x normal value). Other parameters are equivalent to the normal physiological state of RBCs. RBC volume is expressed in % relative to normal isotonic cell volume – applying to all figures in this study. It should be noted that hemolytic volume equals about 160 % of the normal RBC volume.

**Figure 4.**
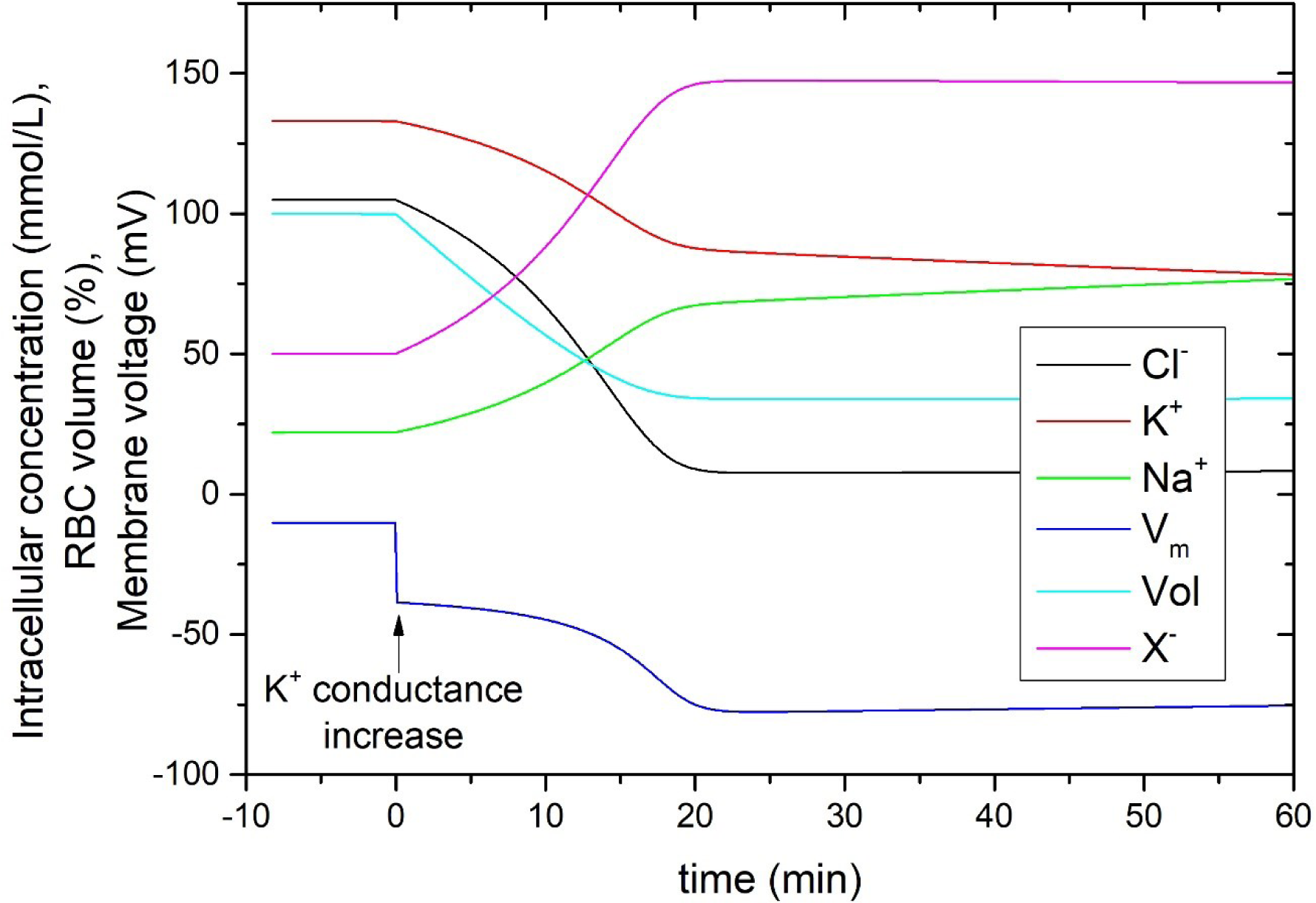
The physiological response (an RVD equivalent) to increased potassium conductance (100 x normal value). Other parameters are equivalent to the normal physiological state of RBCs.

**Figure 5.**
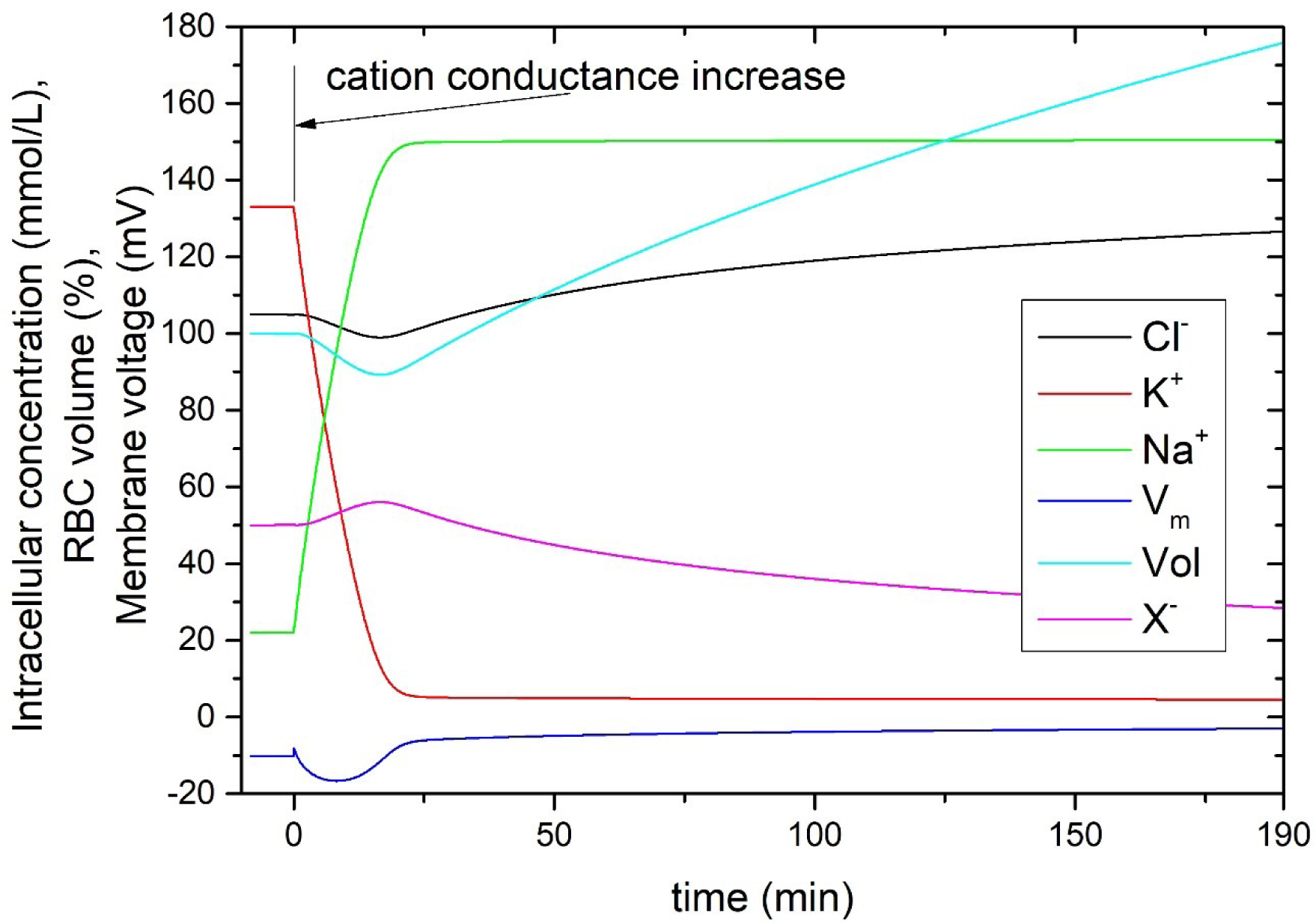
The physiological response to a uniform increase in cationic (Na+ and K+) conductance (100 x normal value). The solution emphasizes that even non-selective ion transport pathways eventually facilitate an RVI response.

Regarding the post-hypertonic lysis, the dynamic evolution of the RBC physiological state shows that sodium-selective pathways facilitate a volume increase, eventually reaching the hemolytic volume (∼160 %). It should be noted that in the case of sodium-selective conductance increase, the gained amount of NaCl resulting in osmotic lysis upon resuspension in an isotonic medium is equal to 50 mmol of NaCl per liter of RBCs. While the current study addresses the hypertonic behavior of human RBCs, the model is solved at isotonic conditions. Within the model and applied approximations (especially invariance of the increased conductance to the level of tonicity), the tonicity at which the numerical simulation is performed does not affect the numerical solutions. A discussion of possible discrepancies is offered later in the section on the limitation of the current study. Regarding the time aspect of the salt loading, it is interesting to know the relationship between the ion flux rate offered by hypertonicity-induced ion transport pathways and the time when the critical hemolytic volume is reached. These findings are shown in Figure 6. The relationship shows a non-linear dependence when a relatively long time is needed for the critical salt loading to complete in the case of a 10-fold sodium conductance increase (3 hours). However, the time quickly falls from hours to tens of minutes upon a 50x conductance increase, with further conductance increase approaching a time interval of about 12 minutes, emphasizing the relatively high rate of such RVI response. While the conductance rates applied in this study may appear arbitrary, it is emphasized in the Discussion section that ion transport pathways in the RBC plasma membrane can facilitate such ion flux rates.

**Figure 6.**
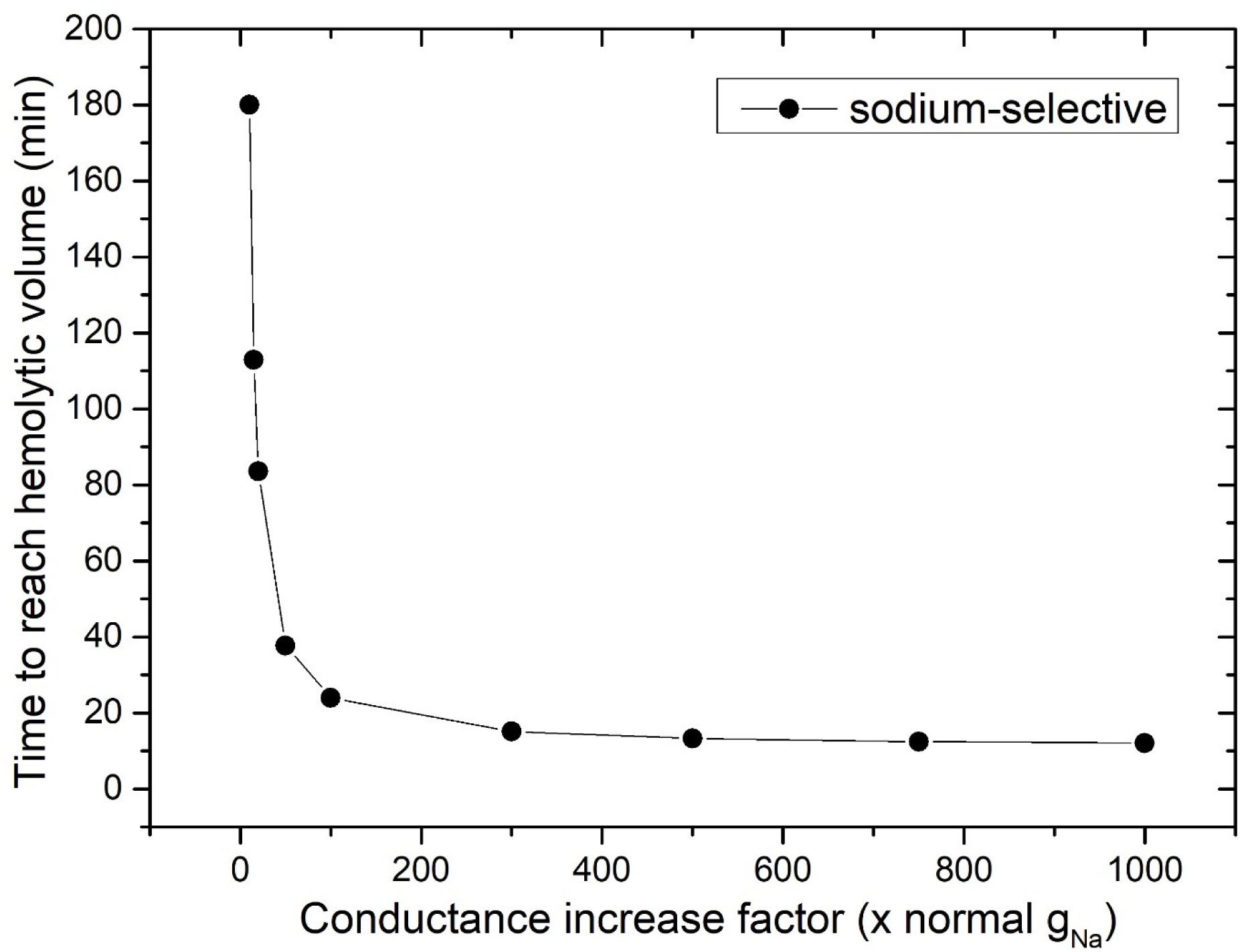
The relationship between the time needed to reach the hemolytic volume and conductance increase.

**Figure 7.**
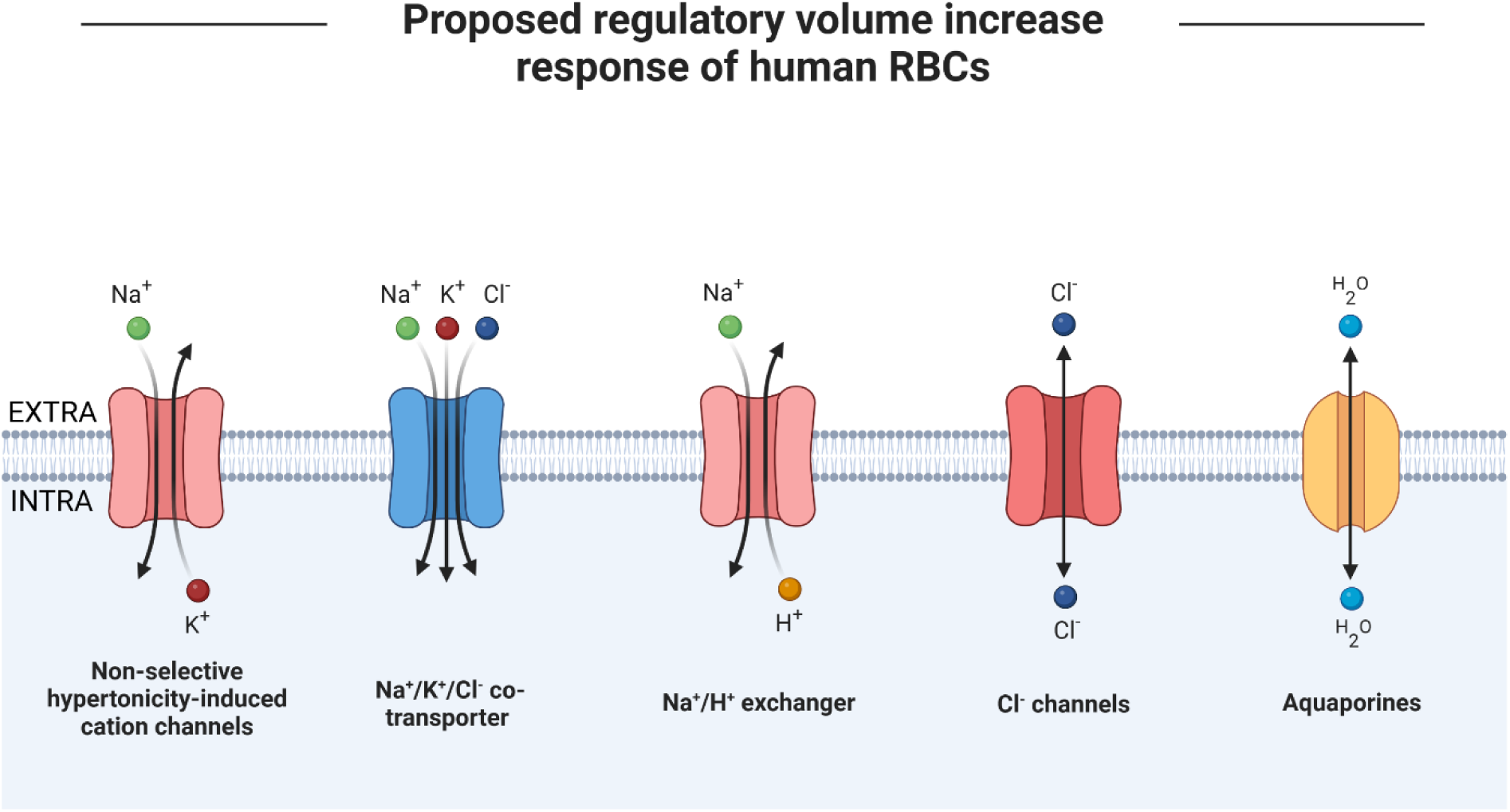
The hypothesized RVI response, including individual ion transport pathways, facilitating the overall loading of NaCl. The picture also emphasizes the passive distribution of chlorine ions and water.

## DISCUSSION

The results of the model based on the pump-leak concept recognizing the dynamic equilibrium between active ion transport and dissipative ion fluxes across plasma membrane support the hypothesized involvement of acute cell volume regulatory mechanisms in RBC osmotic response to hypertonic stress. The model demonstrates how the increased cation permeability explains the cation leaks, post-hypertonic swelling, and post-hypertonic lysis. Specifically, the activation of the ion transport pathways resulting in overall sodium uptake is coupled to membrane depolarization-induced intake of passively distributed chlorine ions, consistent with the expected outcome of the increased sodium permeability.^48^ It is emphasized that the salt loading results from the increased passive membrane permeability to cations and that it occurs via leak pathways without the input of metabolic energy – harvesting the normal electrochemical gradients for cations. While the charge difference method is preferred to solve the dynamic evolution of the RBC physiological state, the role of the applied model is not decisive for the current work, as fundamentally same results are obtained by consulting models based on the GHK approach or its derivatives.^56, 57, 64^ The current study supports the previous suggestions of the osmotic nature of post-hypertonic lysis.^2, 3, 12^ It is argued that post-hypertonic lysis is essentially hypotonic lysis already occurring at isotonic osmolality of the resuspension medium due to an excess of intracellular salt. This is consistent with a work that showed that artificial loading of RBCs with cations leads to a shift in osmotic fragility curves with the onset of the hypotonic lysis occurring at lower tonicity.^65^ The predicted critical amount of excess salt gained by an RVI response, which is 50 mmol of NaCl in the sodium-selective case, roughly matches the sodium influx above which post-hypertonic lysis develops in experiments.^15^ Considering the high permeability of RBC plasma membrane to water, RBCs are expected to be brought into osmotic equilibrium primarily by water fluxes upon resuspension into isotonic medium, inevitably leading to osmotic lysis if the critical number of intracellular salts and associated cell volume are exceeded.

Results are consistent with the outcome of initiating cell volume regulation pathways in proliferation and suicidal form of death (apoptosis), which are associated with isotonic cell swelling and shrinkage, respectively.^40^ The results are also consistent with the pathophysiological states of RBCs caused by altered cation permeability due to the mutations affecting the function of ion transport pathways. Their selectivity and gating might also be affected, potentially resulting in the development of pathological states. Increased sodium permeability leads to sodium uptake and associated higher hydration (a state called “overhydrated stomatocytosis”), characterized by increased hemolysis of circulating RBCs. On the other hand, increased potassium permeability leads to a loss of intracellular potassium and associated dehydration (a state called “xerocytosis”). There is also a pathophysiological state known as “cryohydrocytosis,” which describes a condition with cold-induced cation leak leading to overhydration (for a review of these disorders, see the work of Gallagher^66^). Although these are pathological states, which negatively affect the functionality of the RBCs circulating in the blood, it should be emphasized that the activation of the ion transport pathways under hypertonic conditions, which is understood as an RVI response in this study, should be regarded as a healthy physiological response of RBCs to reduce hypertonicity-related stress by increasing cell volume and maintaining cell functionality. While RVI response is initiated to protect cells, excessive NaCl influx in combination with an abrupt dilution of the bathing medium causes osmotic lysis when critical salt content is exceeded.

Although the normal cation permeability at isotonic conditions is relatively low, the hypertonic activation of cation channels and ion transporters may substantially increase the plasma membrane permeability. There is a certain ambiguity in the assumed cation conductances upon hypothesized activation of ion transport pathways facilitating the RVI response. However, the time aspect of the salt loading predicted in this study is consistent with typical time courses of acute cell volume regulation observed across different kinds of biological cells. ^27, 44, 45^ Specifically to human RBCs, experimental studies also support the involvement of ion transport pathways in the osmotic response of RBCs as the ion flux rates may be high enough to result in considerable KCl loss within ∼15 minutes^67^ (action equivalent to RVD response) or increase of intracellular Na in 30 minutes (RVI).^50^ Furthermore, whole-cell recordings of 68 to 500 pico-siemens cation conductances were reported, ^51, 68, 69^ corresponding to about a 3 to 4 order increase compared to the normal isotonic value.^70^ This experimental observation supports the relatively high rate of predicted NaCl loading occurring only via physiological ion transport pathways (consult Figure 6).

It could be argued that a post-RVI RVD response could save the RBCs from osmotic lysis during resuspension. However, the rates of KCl efflux are not likely to be considerably higher than the predicted NaCl loading; therefore, the osmotic equilibrium is still reached primarily by passive water fluxes due to the high hydraulic conductivity of the RBC plasma membrane. However, the situation might be different when the resuspension into the isotonic medium is performed more gradually, giving time for the RVD response to reduce the number of intracellular salts. This way, RBCs can reach the normal physiological state with the help of ion pumps since normal electrochemical gradients need to be re-established – a process directly coupled to metabolic energy consumption. This prediction is consistent with the observed behavior of RBCs showing lesser hemolysis when RBCs are resuspended into a medium with gradually decreasing tonicity either in isothermal osmotic measurements^18, 19, 71^ or freeze-thaw experiments.^7^ A beneficial effect of slow thawing, equivalent to gradual dilution, was also observed following the slow freezing of human RBCs.^72^

While the current study treats the problem in theory, several correlations with the osmotic behavior of RBCs, either during isothermal osmotic experiments or actual cell cryopreservation, might be provided to validate the proposed hypothesis. The first piece of evidence comes from a study revealing that chlorpromazine – ion channel inhibitor^73, 74^ – reduces post-hypertonic lysis of RBCs.^75^ According to the proposed hypothesis, inhibition of the ion transport pathways should prevent salt loading and associated isotonic cell swelling, thus avoiding post-hypertonic lysis. Another experimental evidence comes from a study by Rudenko and Patelaros, who examined the effect of divalent cations on the post-hypertonic lysis of RBCs.^76, 77^ Although the authors proposed an alternative explanation of the hypertonicity-related RBC behavior when they assumed a formation of post-hypertonic hemolytic pores, it is argued in the current study that their results support the hypothesized involvement of ion transport pathways. As suggested by these authors, the post-hypertonic hemolytic pores were supposed to be affected by divalent cations binding to “activating and inhibitory membrane sites.” However, from the current point of view of modern physiology and membrane biophysics, these membrane pores and sites are likely ion channels and transporters with specific sites sensitive to interaction with potential inhibitors. The authors observed suppression of post-hypertonic lysis of RBCs treated with hypertonic NaCl solution by Ca^2+^ when present in the isotonic resuspension medium. It is known that Ca^2+^ stimulates the potassium-selective Gardos channels.^78–80^ These channels are involved in acute cell volume regulation (RVD response) and the promotion of RBC eryptosis (apoptosis-like cell death) characterized by isotonic cell shrinkage.^81^ It might be expected that the presence of Ca^2+^ in isotonic resuspension medium enhances the K^+^ permeability, affecting the membrane potential and stimulating the loss of intracellular potassium (followed by chlorine) – enhancing the post-RVI RVD, which should have a protective effect against post-hypertonic lysis. Authors also observed an inhibitory effect of Mg^2+^ and Zn^2+^ on post-hypertonic lysis of RBCs, which might be explained by their binding to NMDA receptors – non-selective cation channels possibly involved in the cation leaks, blocking the channel for other cations.^82^

Indirect evidence might be offered by demonstrating the explanatory power of the proposed hypothesis, which may resolve why cells initially suspended in anisotonic solutions differ in susceptibility to subsequent freezing and thawing.^8, 83, 84^ Pegg and Diaper^8^ emphasized anomalous behavior of RBCs initially suspended in 0.6x and 4x isotonic solution differing from those suspended in 1x and 2x isotonic. This might be explained by a more pronounced cell volume regulation associated with a higher level of hypertonicity and hypotonicity of the initial suspension medium. This argument applies to both isothermal osmotic experiments as well as the freezing of cells. While Pegg and coworkers associated the loss of cell viability with cell volume changes, Mazur and colleagues had proposed a different explanation based on the “unfrozen fraction” hypothesis ascribing the deleterious effects to interactions between cells and extracellular ice.^7^ The perspective provided in the current study could offer a unifying viewpoint regarding this debate between Mazur and Pegg and their colleagues. While it is often assumed that the deleterious effect of extracellular ice on cells lies in the adverse effects of mechanical forces acting upon the plasma membrane and disrupting it structurally,^7, 85, 86^ it is also possible that these forces induce cation leaks since many ion transport pathways exhibit some mechanosensitivity,^87^ while it was also observed that mechanical force induces cation leaks of human RBCs.^88^

As mentioned before, only Muldrew considered the involvement of ion channels in the salt loading of hypertonicity-treated human RBCs. With his “salting in” hypothesis of post-hypertonic lysis, Muldrew suggested that sodium ions enter the cytoplasm via ion channels that open during hypertonic conditions and that these ions replace the potassium ions in cytoplasmic solution, which bind to salted- in intracellular proteins upon concentration in cytoplasm.^20^ The author further hypothesized that during resuspension to isotonic medium, the intracellular proteins are salted-out of the solution, releasing the bound potassium and thus increasing the number of osmotically active solutes in the cell cytoplasm, leading to osmotic lysis if critical concentration and associated cell volume are exceeded. Compared to Muldrew’s hypothesis, this work provides a more advanced physiological perspective on the possible involvement of ion channels, including consideration of membrane potential and discussion of different types of ion transport pathways possibly involved in the RVI response (for the detailed discussion, see the section of the text on identification of the ion transport pathways involved in the osmotic response). While the salting-in hypothesis did not consider the effect of membrane voltage, the theory presented in this paper includes the calculation of membrane potential – the overall thermodynamic driving force for ions is given by electrochemical gradients. Also, based on the perspective of cell volume regulation, the assumption of potassium binding to intracellular proteins is avoided in the current hypothesis. The cation channels possibly involved in the cation leaks are also specified to the class of hypertonicity-induced cation channels, allowing their possible identification and targeting to minimize the cation leaks and reduce post-hypertonic lysis of RBCs.

Regarding alternative explanations, Muldrew^20^ also mentioned an alternative explanation to post-hypertonic lysis, based on the observation of changes in plasma membrane surface area of protoplasts exposed to hypertonicity,^89^ claiming that reduction of the surface area could induce lysis in isotonic resuspension medium. This idea is, however, at least in part inconsistent with the observed behavior of human RBCs, which show swelling beyond normal cell volume upon resuspension.^77^ Another alternative explanation could be provided by consulting the osmotic rupture hypothesis proposed by Muldrew and McGann,^90, 91^ which claims the osmotically-induced water fluxes to be detrimental, causing membrane damage. However, the author of the current study is unaware of any report concerning human RBCs which would support this explanation. Some evidence against the involvement of this effect in the post-hypertonic lysis of human RBCs comes from a study on the post-hypertonic lysis of PC-3 cells,^11^ which demonstrated that the rate of water fluxes does not have a significant effect and that post-hypertonic lysis is determined by the duration of exposure to hypertonicity and its level. The current hypothesis also represents an alternative explanation to any idea considering the formation of membrane lesions or any even reversible loss of plasma membrane integrity. In contrast with these hypotheses, the involvement of cell volume regulation mechanisms easily explains the absence of simultaneous loss of hemoglobin or sucrose gains^15^ (when present in an extracellular solution) since ion transport mechanisms are not expected to be permeable to hemoglobin or sucrose.

### Limitations of the model

Firstly, some discrepancies between predicted and actual RBC behavior may arise from differing activity coefficients of ion species across the plasma membrane and the increased osmotic activity of cytoplasmic hemoglobin as its concentration in intracellular fluid rises. Since the basic physiological model proposed in this study already explains the salt loading and post-hypertonic lysis without their consideration, the focus is placed on fundamental governing physiological principles, and their account is omitted in the current study. Secondly, the fact that the outlined model is solved for isotonic conditions also represents a certain limitation of the current study. The formulation of the model is not limited to isotonic conditions; in principle, it is valid for any level of tonicity and associated cell hydration. Within the approximation of constant ion conductances, the model represents the exact solution to the hypertonic experiment, irrespective of the tonicity at which the calculations are performed. Some discrepancies might be introduced with the assumption of constant conductances, which may be subject to change upon salt concentration increase. Nevertheless, it is emphasized that this limitation affects only the time aspect of the salt loading, not the endpoint (equilibrium state). It is also argued that the error introduced by this approximation has not decisive role neither in the time aspect of salt loading, as the conductance increase is presumably affected mainly by the hypertonic activation of the ion transport pathways that are almost inactive at normal isotonic conditions. Also, as the applied model predicts, the time needed for accumulating a critical amount of NaCl saturates above 100x increase of sodium conductance (see Figure 6) implies that possible discrepancy should not be considerable. More sophisticated physiological models might be applied in the future, including consideration of pH changes, calcium ions, which have a role in RVD initiation, and the interplay between individual ion pathways might also be involved. It appears to the author of this study that these effects would unnecessarily complicate the picture for the current purpose of this work. Therefore, the consideration of these effects is omitted.

### Identification of the ion transport pathways involved in the osmotic response

The simplest possible explanation for the sodium intake occurring under hypertonic conditions is offered by hypothesizing that HICCs and other ion transport pathways involved in the RVI have high selectivity for sodium. While sodium-selective channels involved in RVI were identified in other cells,^48^ such channels were not identified in human RBCs. Their HICCs appear to be non-selective to cations.^49^ Even though the overall response is consistent with the activation of strongly sodium-selective ion transport pathways, the actual ion movements might be more complex. Theoretically, the same result would be obtained if the HICCs were non-selective but would perform in concert with other ion transport mechanisms, which transport potassium back into the cells. One obvious candidate is the sodium-potassium pump. However, the capacity of these pumps in human RBCs is limited, implying no significant role of the pump in the osmotic response. In addition, the pump also extrudes the sodium out of cells, which would be counterproductive from the perspective of RVI response. On the other hand, ion transporters could facilitate the intake of lost potassium. A candidate for such action is NKCC transporter,^92–94^ co-transporting Na^+^, K^+^, and 2 Cl^-^ ions, identified in human RBCs’ RVI response.^95, 96^ Note that these transporters are classified as secondary active ion transporters, driven by electrochemical gradients, not by the consumption of ATP. Although the K^+^ ions are transported back into the cell against their electrochemical gradient, the cotransport is associated with an overall negative change of Gibbs energy as Na^+^ and Cl^-^ move down their electrochemical gradients under depolarizing conditions, making it a leak pathway. A further contribution to the salt loading might be provided by activation of the Na^+^/H^+^ exchanger (increasing the overall sodium-selectivity), which was also identified as an element of the RVI response of human RBCs.^94, 97^

Regarding the identification of the HICCs involved in the RVI response of human RBCs, a study revealing that mechanical stress applied to RBCs induces cation leaks in isotonic and anisotonic media should be mentioned.^88^ This observation supports the mechano-sensitivity of cation channels hypothesized to be involved in cation leaks and post-hypertonic lysis. Many ion channels exhibit mechanosensitivity.^87^ Specifically to human RBCs, this includes PIEZO1 channels,^87, 98^ transient receptor potential channels TRPC6, ^99^ and TRPV2. ^100–102^ Based on the works of Rudenko and Patelaros discussed above, it might be argued that NMDA receptors^103^ could also be involved in hypertonicity-related cation leaks, and there is also a report for non-selective cation channels in the RBC membrane activated by membrane depolarization.^52, 104, 105^

## CRYOBIOLOGICAL SIGNIFICANCE

Only a few studies have considered the involvement of cell volume regulation mechanisms in cryobiologically relevant processes. Since cells subject to cryopreservation are also exposed to hypertonic conditions – due to the freeze-concentration process as extracellular ice forms – the importance of the proposed hypothesis for cryopreservation and freezing injury of cells is evident. In the past, Peckys and Mazur doubted the possible involvement of cell volume regulation mechanisms in the osmotic response of cells during freezing when they assumed an inhibitory effect of reduced temperature on ion transport rates.^21^ However, they considered only the participation of the active ion transport, which is directly coupled to the consumption of metabolic energy. However, the most effective acute cell volume regulation elements are leak pathways allowing ions to move down their electrochemical gradients. It is also emphasized in this study that electrochemical gradients for cations are always present in normal healthy cells, giving the capacity to cells to perform acute cell volume regulation. The passive cation permeability might be easily increased by triggering the dissipative ion transport pathways, which are normally present in the RBC plasma membrane but inactive at normal isotonic conditions. Since ion channels allow passive dissipative ion fluxes without the need for input of metabolic energy, the effect of reduced temperature is also not expected to be dramatic. As long as ion channels remain open, ion fluxes are expected to occur, although some decrease in ion flux rates might be expected due to temperature-reduced ionic mobility. It should be, however, noted that some ion channels are activated by reduced temperature;^106^ therefore, the actual temperature dependence of cation permeability might be a non-monotonous function of temperature. That might be relevant also to RBCs as witnessed by an experimental study by Stewart, which showed the generally expected decrease of ion flux rate with reduced temperature, reaching a local minimum and increasing with further temperature reduction below 12°C.^107^

### On (cryo)protective action of cryoprotective agents

The protective action of the cryoprotectants against slow-freezing injury has often been ascribed to the non-specific reduction of salt concentrations. Regarding salt loading, it was shown that dimethyl sulfoxide, a common cryoprotectant, reduces the cation leaks and postpones their onset to higher salt concentrations when present in the hypertonic medium,^13^ suppressing the post-hypertonic lysis. The same effect had dimethyl sulfoxide on hypertonicity-treated Chinese hamster ovary cells. ^108^ From the physiological perspective, dimethyl sulfoxide, glycerol, and similar permeant additives reduce salt concentrations on both sides of the plasma membrane, thus not altering the ion electrochemical gradients across the plasma membrane. Within the logic of the formulated hypothesis, a possible mechanism of cryoprotective action could be associated with an inhibitory effect of cryoprotectants on ion channels. Indeed, there is evidence that dimethyl sulfoxide inhibits the non-selective cation channels of human erythrocytes.^68^ Inhibition of these ion channels would decrease the extent of cation leaks and thus reduce the post-hypertonic lysis. The mechanism of the inhibitory effect of dimethyl sulfoxide on the non-selective cation channels has yet to be elucidated. It might be speculated that dimethyl sulfoxide could shift the structural conformation of ion channels towards the conformation representing the closed state due to molecular interactions with proteins forming the ion channels, water, or by alteration of the physical properties of the lipid bilayer and thus affecting the protein-lipid interactions. However, while dimethyl sulfoxide was found to inhibit the cation fluxes, glycerol did not affect them.^68^ Since both additives offer cryoprotection to RBCs, these results suggest that the protective action of these agents might also be facilitated by their non-specific effect on the stimuli triggering the ion channels. The non-selective cation channels of human RBCs were shown to be sensitive to cell volume and internal and external Cl^-^ concentration.^49^ Both these triggers are affected upon loading cells with permeant cryoprotectants. For instance, at a given osmotic concentration of a bathing solution, the intracellular concentrations of ions are reduced due to the presence of permeant additive as well as the absolute cell volume is larger since these additives increase the overall number of osmotically active solutes in the cytoplasm – an outcome consistent with the aim of RVI response. These considerations provide a mechanism for non-specific protection by permeant (cryo)protectants.

While permeant cryoprotectants offer protection against post-hypertonic lysis, non-permeants do not.^18, 19^ RBCs treated with hypertonic non-electrolyte solutions are subject to cell dehydration of the same magnitude as those exposed to hypertonic electrolyte solutions of the same osmolality. Considering the minor content of salts in these solutions, the capacity to perform RVI is limited due to the minor content of extracellular NaCl. In addition, the dilution of extracellular salts by non-permeant additives disturbs the electrochemical gradients of all ions, inducing an outward electrochemical gradient for passively distributed chlorine ions - already at isotonic conditions. Overall, KCl loss might be expected under such conditions, exhibiting a considerable increase in the flux rate as ionic strength reaches very low values. ^53^ At first, it seems impossible for cells to undergo osmotic lysis upon resuspension into isotonic (either salt or again non-electrolyte) medium since the intracellular number of salts should be lower than normally – resulting in isotonic shrinkage of RBCs. In this respect, an interesting question arises when addressing the resuspension phase of such osmotic experiments: does the salt loading occur only under hypertonic conditions or also at resuspension to the isotonic medium? It is not easy to judge this from the cited experimental studies. While the cell shrinkage vanishes upon resuspension into an isotonic medium because of associated cell swelling, it is unclear whether the ion transport pathways inactivate immediately. If they remain activated, there would be a massive inward driving force for NaCl entry into the cells resuspended in an isotonic salt solution following treatment with a hypertonic non-electrolyte solution. A study addressing the protective effect of chlorpromazine speaks in favor of this suggestion, as the protective effect was most pronounced when the channel inhibitor was present in the resuspension medium.^109^ Further evidence comes from a study demonstrating that an RBC voltage-gated non-selective cation channel, possibly involved in the RVI response, exhibits a hysteretic dependence on membrane voltage – remaining open at conditions at which it was closed before activation.^110^ The findings of Christmann et al. also support this idea when the freezing was observed to stimulate the post-cryo RVI response of HepG2 and HeLa cells.^24^ This could also explain the complex nature of divalentś effect on post-hypertonic lysis of RBCs treated with hypertonic non-electrolyte solutions in the studies by Rudenko and Patelaros.^76, 77^

While the current study suggests the osmotic nature of the post-hypertonic lysis, which appears to be the dominant form of osmotic injury of human RBCs, the KCl loss stimulated by non-permeant additives might induce other forms of injury as well – for example, eryptosis. Therefore, from the perspective of osmotic injury, the non-permeant additives induce stress rather than offer protection. The only protective aspect of the non-permeants (regarding post-hypertonic lysis) seems to lie in their ability to increase the osmolality of the resuspension media, avoiding post-hypertonic lysis – simply because RBCs do not reach the critical hemolytic volume due to the hypertonic nature of the resuspension medium. That might allow RBCs to lose the accumulated salt and reach the normal physiological state with the help of active ion transport and RVD mechanisms. However, beyond post-hypertonic lysis, other forms of cryoinjury are suppressed by non-permeant additives, such as ice formation or eutectic salt crystallization, explaining their utility in cryopreservation, especially in combination with permeant cryoprotectants or in alternative approaches reducing the osmotic injury in other ways (see the discussion on pre-freeze induced cell dehydration cryopreservation protocols in the following section).

### Implications for the design of RBC cryopreservation protocols

The presented hypothesis has several implications for the design of the cryopreservation protocols. The most important implication is the possibility of reducing the post-hypertonic lysis and slow-freezing injury of RBCs by introducing inhibitors of specific ion transport pathways into the hypertonic (or cell freezing) medium. The hypothesis predicts that adding ion channel blockers into the hypertonic or resuspension medium will suppress the cation leaks and thus reduce the extent of post-hypertonic lysis of human RBCs. There is already a study demonstrating the protective effect of chlorpromazine – an ion channel blocker – against post-hypertonic lysis of RBCs.^75^ Considering the cell volume regulation perspective, other ion channel blockers specifically targeting the HICCs, such as amiloride, flufenamate, or Gd^3+^,^27^ could protect RBCs from post-hypertonic lysis. This prediction is consistent with the observation by Duranton et al., who showed inhibition of cation conductance by amiloride, resulting in the inhibition of associated hemolysis.^51^ Introducing inhibitors targeting the ion transport pathways involved in RVD could also improve post-thaw cell recovery since these are also initiators of apoptosis, which has been identified in cells’ post-thaw viability evaluation. However, it should be emphasized that other forms of cryoinjury – such as ice formation or eutectic crystallization,^7, 111, 112^ are unlikely to be suppressed by these inhibitors, typically introduced at sub-millimolar concentrations. For example, the concentration of dimethyl sulfoxide needs to be at least 2 vol % (∼260 mmol/L) in isotonic NaCl-based cell media to suppress the eutectic crystallization of the NaCl entirely.^113^ Therefore, cryopreservation protocols introducing ion channel blockers would likely still require a common cryoprotectant, although perhaps at a reduced concentration than conventional protocols, often relying on 10 vol % dimethyl sulfoxide or even higher concentrations of glycerol in RBC cryopreservation. Another implication of the proposed hypothesis is that supplementing the sodium with other plasma membrane impermeable cations, such as choline (see the following section of the text addressing the choline-based freezing media in oocyte cryopreservation), could reduce the post-hypertonic lysis considerably.

The post-thaw cell viability could also be increased by applying freezing protocols relying on a high cooling rate, recognizing the time aspect of post-hypertonic lysis and slow-freezing injury. However, rapid freezing protocols are dangerous because of insufficient cell dehydration and associated detrimental intracellular ice formation. In this respect, cryopreservation protocols exploiting the pre-freeze-induced cell dehydration may represent a promising approach for designing future cryoprotective mixtures and protocols, possibly with a significantly reduced content of typically applied permeant cryoprotectants. In these protocols, the applied high cooling rate minimizes the time-dependent slow freezing injury, and the controlled pre-freeze cell dehydration allows freezing without the risk of intracellular ice formation. Recent reports show the successful application of pre-freeze-induced cell dehydration in the cryopreservation of fibroblasts, RBCs, and keratinocytes.^114–116^ It should also be noted that cryopreservation protocols for RBCs were reported in the past, ^117, 118^ relying only on extracellular cryoprotectants and rapid freezing. The success of these protocols might also be rationalized by considering the pre-freeze-induced cell dehydration.

### Broader cryobiological significance

Besides the RBC-related phenomena, the formulated hypothesis might be relevant also to other cell types since cell physiology described in this paper, including the mechanisms of acute cell volume regulation, applies practically to all types of cells (for a picture of which cell volume regulation mechanisms are present in which cell type, an extensive summary covering a broad range of cells is offered in a review by Lang et al.^38^). The proposed hypothesis also explains the deviations of the cell osmotic response from the expected behavior when unexpected volume changes or the inability of cells to return to the initial volume were observed. ^29–31^ Therefore, cation leaks and associated post-hypertonic lysis appear to be a general mechanism of osmotic cell injury, which might be expected to be most pronounced in slow-freezing cryopreservation protocols. The current hypothesis also explains the unexpected freezing-induced swelling of human pulmonary microvascular endothelial cells,^119^ which occurred instead of the expected cell shrinkage induced by the freeze-concentration induced by ice formation. Note that endothelial cells are equipped with a HICC called ENaC with high selectivity for Na^+^ ions,^120^ implying a more efficient RVI response since the net uptake of ions and associated water influx is more significant than in the case of non-selective channels. Note that a study on the osmotic injury of the PC-3 cells (exhibiting post-hypertonic lysis) showed that some of the cells regained their size already in the hypertonic phase of the experiment.^11^ Another observation, which might be explained, is the insensitivity of mouse oocytes,^121, 122^ human oocytes,^123^ or ovarian tissue^124^ to slow freezing injury when choline ions replace the sodium ions in the cell freezing medium. As suggested by a study on Ehrlich-Lettre-ascites cells, HICCs allow permeation of Na^+^, K^+^, and Li^+^ but are impermeable to choline^+^ or N-methyl-d-glucamine^+^.^125^

While the current study focuses on post-hypertonic lysis, other forms of osmotic injury have been observed during osmotic experiments with RBCs. Hypertonicity-treated cells were observed to be sensitive to abrupt changes in temperature or mechanical shock associated with their centrifugation.^2, 17, 126, 127^ Although these phenomena are beyond the scope of this work, ion transport pathways might also be involved in those situations. As mentioned before, the cation leaks are also stimulated by mechanical force. The centrifugal forces might not be high enough under isotonic conditions to induce cation leaks. However, at higher tonicities, these forces may cause cation leaks at lower onsetting tonicity – an inference consistent with the study by Shpakova et al.,^88^ which showed mechanically induced cation leaks and post-hypertonic lysis at lower tonicities when compared to mechanically unstimulated RBCs. The detrimental aspect of temperature is less obviously linked to the activation of ion transport pathways. It could be suggested that the presence of cation channels stimulated by reduced temperature could induce cation leaks. However, this is somewhat speculative at the current moment.

## SUMMARY

It is concluded that quantitative analysis provided by solving a simplified physiological model supports the hypothesized involvement of cell volume regulation mechanisms in the osmotic response of hypertonicity-treated human RBCs. The current study avoids the commonly applied assumption of plasma membrane impermeability to ions, which is fundamentally inconsistent with current cell physiology and is problematic, especially at anisotonic conditions, when cell volume regulation might be initiated. It is shown that the RBC plasma membrane contains ion transport pathways, which facilitate regulatory volume increase response, and it is also shown that their hypertonic activation might facilitate a several-order increase in cation permeability without loss of plasma membrane integrity. It is also emphasized that the most efficient elements of acute cell volume regulation are leak pathways – ion channels and transporters – facilitating passive ion dissipative fluxes, exploiting the normal electrochemical gradients for cations. The current hypothesis claims that post-hypertonic lysis is essentially a hypotonic lysis occurring already at isotonic osmolality due to accumulated salt. The mechanism of protective action of common permeant cryoprotectants might be primarily ascribed to non-specific inhibition of triggers initiating an RVI response – cell shrinkage and increased ion concentrations. Perhaps the most important practical implication of the proposed hypothesis is the possibility of avoiding post-hypertonic lysis and thus slow-freezing injury by introducing inhibitors specifically targeting ion transport pathways – a prediction already consistent with few experimental studies. A broader significance of the outlined hypothesis is evident since cell volume regulation mechanisms are present in many biological cells, suggesting a general mechanism of osmotic injury or “solution effects” injury associated with cell cryopreservation. A high explanatory power of the proposed hypothesis is also demonstrated by explaining the complex behavior of RBCs either in isothermal osmotic experiments or actual cryopreservation. Furthermore, the hypothesis also explains the non-ideal osmotic behavior of other cells, including post-hypertonic lysis, cation leaks, unexpected freezing-induced cell swelling, and the reduced “solution effects” injury associated with sodium replacement by impermeable choline in the cell medium during slow freezing cryopreservation. Also, an alternative mechanism for cryoinjury imposed by ice formation was proposed by considering the mechanical activation of cation channels involved in cation leaks. Overall, the current work clearly emphasizes the utility of the physiological perspective in cryobiological considerations, two fields that have had historically rather low overlap.

## References

1. Takei T. Über die Analyse einer Volumkurve von Blutkörperchen in hypertonischen Lösungen, welche zugleich die Differenzerung von osmotischen und kolloidehemischen Volumänderungen ermöglicht. Biochem Z. 1921;123:104–127.

2. Lovelock JE. The haemolysis of human red blood-cells by freezing and thawing. Biochim Biophys Acta. 1953;10:414–426. doi:10.1016/0006-3002(53)90273-X

3. Zade-Oppen AMM. Posthypertonic Hemolysis in Sodium Chloride Systems. Acta Physiol Scand. 1968;73(3):341–364. doi:10.1111/j.1748-1716.1968.tb04113.x

4. Zade-Oppen AMM. Posthypertonic hemolysis in a sucrose system. Experientia. 1970;26(10):1087–1088. doi:10.1007/BF02112690

5. Zhurova M, Lusianti RE, Higgins AZ, Acker JP. Osmotic tolerance limits of red blood cells from umbilical cord blood. Cryobiology. 2014;69(1):48–54. doi:10.1016/j.cryobiol.2014.05.001

6. Semionova EA, Yershova NA, Yershov SS, Orlova N V., Shpakova NM. Peculiarities of posthypertonic lysis in erythrocytes of several mammals. Problems of Cryobiology and Cryomedicine. 2016;26(1):73–83. doi:10.15407/cryo26.01.073

7. Mazur P, Cole KW. Roles of unfrozen fraction, salt concentration, and changes in cell volume in the survival of frozen human erythrocytes. Cryobiology. 1989;26(1):1–29. doi:10.1016/0011-2240(89)90030-8

8. Pegg DE, Diaper MP. The effect of initial tonicity on freeze/thaw injury to human red cells suspended in solutions of sodium chloride. Cryobiology. 1991;28(1):18–35. doi:10.1016/0011-2240(91)90004-8

9. Armitage WJ, Mazur P. Osmotic tolerance of human granulocytes. American Journal of Physiology-Cell Physiology. 1984;247(5):C373–C381. doi:10.1152/ajpcell.1984.247.5.C373

10. Gao DY, Ashworth E, Watson PF, Kleinhans FW, Mazur P, Critser JK. Hyperosmotic Tolerance of Human Spermatozoa: Separate Effects of Glycerol, Sodium Chloride, and Sucrose on Spermolysis1. Biol Reprod. 1993;49(1):112–123. doi:10.1095/biolreprod49.1.112

11. Zawlodzka S, Takamatsu H. Osmotic injury of PC-3 cells by hypertonic NaCl solutions at temperatures above 0°C. Cryobiology. 2005;50(1):58–70. doi:10.1016/j.cryobiol.2004.10.004

12. Söderström. N. Hemolysis by Hypertonic Solutions of Neutral Salts. Acta Physiol Scand. 1944;7(1):56–68. doi:10.1111/j.1748-1716.1944.tb03013.x

13. Farrant J. Human red cells under hypertonic conditions; A model system for investigating freezing damage. Cryobiology. 1972;9(2):131–136. doi:10.1016/0011-2240(72)90020-X

14. Farrant J, Woolgar AE. Human red cells under hypertonic conditions; A model system for investigating freezing damage. Cryobiology. 1972;9(1):9–15. doi:10.1016/0011-2240(72)90003-X

15. Farrant J, Woolgar AE. Human red cells under hypertonic conditions; A model system for investigating freezing damage. Cryobiology. 1972;9(1):16–21. doi:10.1016/0011-2240(72)90004-1

16. Meryman HT. Modified Model for the Mechanism of Freezing Injury in Erythrocytes. Nature. 1968;218(5139):333–336. doi:10.1038/218333a0

17. Meryman HT. Osmotic stress as a mechanism of freezing injury. Cryobiology. 1971;8(5):489–500. doi:10.1016/0011-2240(71)90040-X

18. Woolgar AE. Hemolysis of human red blood cells by freezing and thawing in solutions containing polyvinylpyrrolidone: Relationship with posthypertonic hemolysis and solute movements. Cryobiology. 1974;11(1):52–59. doi:10.1016/0011-2240(74)90038-8

19. Woolgar AE. Hemolysis of human red blood cells by freezing and thawing in solutions containing sucrose: Relationship with posthypertonic hemolysis and solute movements. Cryobiology. 1974;11(1):44–51. doi:10.1016/0011-2240(74)90037-6

20. Muldrew K. The salting-in hypothesis of post-hypertonic lysis. Cryobiology. 2008;57(3):251–256. doi:10.1016/j.cryobiol.2008.09.007

21. Peckys D, Mazur P. Regulatory volume decrease in COS-7 cells at 22°C and its influence on the Boyle van’t Hoff relation and the determination of the osmotically inactive volume. Cryobiology. 2012;65(1):74–78. doi:10.1016/j.cryobiol.2012.03.008

22. Lang F. Mechanisms and Significance of Cell Volume Regulation. J Am Coll Nutr. 2007;26(January 2015):613S–623S. doi:10.1080/07315724.2007.10719667

23. Wehner F. Cell Volume-Regulated Cation Channels. In: Mechanisms and Significance of Cell Volume Regulation. Vol 152. ; 2006:25–53.

24. Christmann J, Azer L, Dörr D, Fuhr GR, Bastiaens PIH, Wehner F. Adaptive responses of cell hydration to a low temperature arrest. Journal of Physiology. 2016;594(6):1663–1676. doi:10.1113/JP271245

25. Okada Y. Channelling frozen cells to survival after thawing: Opening the door to cryo-physiology. Journal of Physiology. 2016;594(6):1523–1524. doi:10.1113/JP271842

26. Fraser JA, Huang CLH. A quantitative analysis of cell volume and resting potential determination and regulation in excitable cells. Journal of Physiology. 2004;559(2):459–478. doi:10.1113/jphysiol.2004.065706

27. Hoffmann EK, Lambert IH, Pedersen SF. Physiology of cell volume regulation in vertebrates. Physiol Rev. 2009;89(1):193–277. doi:10.1152/physrev.00037.2007

28. Strange K. Cellular and Molecular Physiology of Cell Volume Regulation. (Strange K, ed.). CRC Press; 2020. doi:10.1201/9780367812140

29. Azam I, Olver D, Benson J. Cryobiological Implications and Measurement of Osmotic Behavior of Human Hepatoma HepG2 Cell. Cryobiology. 2021;103:167. doi:10.1016/j.cryobiol.2021.11.040

30. Casula E, Asuni GP, Sogos V, Fadda S, Delogu F, Cincotti A. Osmotic behaviour of human mesenchymal stem cells: Implications for cryopreservation. PLoS One. 2017;12(9):1–21. doi:10.1371/journal.pone.0184180

31. Casula E, Asuni GP, Sogos V, Cincotti A. HMSCs from UCB: Isolation, characterization and determination of osmotic properties for optimal cryopreservation. Chem Eng Trans. 2015;43(2004):265–270. doi:10.3303/CET1543045

32. Traversari G, Cincotti A. Insights into the model of non-perfect osmometer cells for cryopreservation: A parametric sweep analysis. Cryobiology. 2021;100(November 2020):193–211. doi:10.1016/j.cryobiol.2020.11.013

33. Casula E, Traversari G, Fadda S, Klymenko O V., Kontoravdi C, Cincotti A. Modelling the osmotic behaviour of human mesenchymal stem cells. Biochem Eng J. 2019;151:107296. doi:10.1016/j.bej.2019.107296

34. Olver DJ, Azam I, Benson JD. Meta-analysis and experimental re-evaluation of the Boyle van ‘t Hoff relation with osmoregulation modelled by linear elastic principles and ion-osmolyte leakage. bioRxiv. Published online January 1, 2022:2022.03.05.483010. doi:10.1101/2022.03.05.483010

35. Ioav Cabantchik Z, Knauf PA, Rothstein A. The anion transport system of the red blood cell The role of membrane protein evaluated by the use of ‘probes.’ Biochimica et Biophysica Acta (BBA) - Reviews on Biomembranes. 1978;515(3):239–302. doi:10.1016/0304-4157(78)90016-3

36. Ataullakhanov FI, Korunova NO, Spiridonov IS, Pivovarov IO, Kalyagina N V., Martinov M V. How erythrocyte volume is regulated, or what mathematical models can and cannot do for biology. Biochem (Mosc) Suppl Ser A Membr Cell Biol. 2009;3(2):101–115. doi:10.1134/S1990747809020019

37. Scheer A wilhelm, Kruppke H, Heib R. Red Cell Membrane Transport in Health and Disease. (Bernhardt I, Ellory JC, eds.). Springer Berlin Heidelberg; 2003. doi:10.1007/978-3-662-05181-8

38. Lang F, Busch GL, Völkl H. The Diversity of Volume Regulatory Mechanisms. Cellular Physiology and Biochemistry. 1998;8(1-2):1–45. doi:10.1159/000016269

39. Lang F, Busch GL, Ritter M, et al. Functional significance of cell volume regulatory mechanisms. Physiol Rev. 1998;78(1):247–306. doi:10.1152/physrev.1998.78.1.247

40. Lang F, Föller M, Lang KS, et al. Ion channels in cell proliferation and apoptotic cell death. Journal of Membrane Biology. 2005;205(3):147–157. doi:10.1007/s00232-005-0780-5

41. Bortner CD, Cidlowski JA. Apoptotic volume decrease and the incredible shrinking cell. Cell Death Differ. 2002;9(12):1307–1310. doi:10.1038/sj.cdd.4401126

42. Okada Y, Maeno E. Apoptosis, cell volume regulation and volume-regulatory chloride channels. Comparative Biochemistry and Physiology - A Molecular and Integrative Physiology. 2001;130(3):377–383. doi:10.1016/S1095-6433(01)00424-X

43. García-Pérez A, Ferraris JD. Aldose Reductase Gene Expression and Osmoregulation in Mammalian Renal Cells. In: Strange K, ed. Cellular and Molecular Physiology of Cell Volume Regulation. CRC Press; 1994:373–382. doi:10.1201/9780367812140

44. Cacace VI, Finkelsteyn AG, Tasso LM, Kusnier CF, Gomez KA, Fischbarg J. Regulatory volume increase and regulatory volume decrease responses in HL-1 atrial myocytes. Cellular Physiology and Biochemistry. 2014;33(6):1745–1757. doi:10.1159/000362955

45. Koos B, Christmann J, Plettenberg S, et al. Hypertonicity-induced cation channels in HepG2 cells: architecture and role in proliferation vs. apoptosis. Journal of Physiology. 2018;596(7):1227–1241. doi:10.1113/JP275827

46. Kaestner L. Cation Channels in Erythrocytes - Historical and Future Perspective. Open Biol J. 2011;4(1):27–34. doi:10.2174/1874196701104010027

47. Thomas SLY, Bouyer G, Cueff A, Egée S, Glogowska E, Ollivaux C. Ion channels in human red blood cell membrane: Actors or relics? Blood Cells Mol Dis. 2011;46(4):261–265. doi:10.1016/j.bcmd.2011.02.007

48. Wehner F, Bondarava M, Ter Veld F, Endl E, Nürnberger HR, Li T. Hypertonicity-induced cation channels. Acta Physiologica. 2006;187(1-2):21–25. doi:10.1111/j.1748-1716.2006.01561.x

49. Huber SM, Gamper N, Lang F. Chloride conductance and volume-regulatory nonselective cation conductance in human red blood cell ghosts. Pflugers Arch. 2001;441(4):551–558. doi:10.1007/s004240000456

50. Monedero Alonso D, Pérès L, Hatem A, Bouyer G, Egée S. The Chloride Conductance Inhibitor NS3623 Enhances the Activity of a Non-selective Cation Channel in Hyperpolarizing Conditions. Front Physiol. 2021;12(October):1–12. doi:10.3389/fphys.2021.743094

51. Duranton C, Huber SM, Lang F. Oxidation induces a Cl--dependent cation conductance in human red blood cells. Journal of Physiology. 2002;539(3):847–855. doi:10.1113/jphysiol.2001.013040

52. Christophersen P, Bennekou P. Evidence for a voltage-gated, non-selective cation channel in the human red cell membrane. BBA - Biomembranes. 1991;1065(1):103–106. doi:10.1016/0005-2736(91)90017-3

53. LaCelle PL, Rothsteto A. The passive permeability of the red blood cell in cations. J Gen Physiol. 1966;50(1):171–188. doi:10.1085/jgp.50.1.171

54. Goldman DE. POTENTIAL, IMPEDANCE, AND RECTIFICATION IN MEMBRANES. Journal of General Physiology. 1943;27(1):37–60. doi:10.1085/jgp.27.1.37

55. Hodgkin AL, Katz B. The effect of sodium ions on the electrical activity of the giant axon of the squid. J Physiol. 1949;108(1):37–77. doi:10.1113/jphysiol.1949.sp004310

56. Mullins LJ, Noda K. The Influence of Sodium-Free Solutions on the Membrane Potential of Frog Muscle Fibers. Journal of General Physiology. 1963;47(1):117–132. doi:10.1085/jgp.47.1.117

57. Armstrong CM. The Na/K pump, Cl ion, and osmotic stabilization of cells. Proceedings of the National Academy of Sciences. 2003;100(10):6257–6262. doi:10.1073/pnas.0931278100

58. Fraser JA, Huang CLH. Quantitative techniques for steady-state calculation and dynamic integrated modelling of membrane potential and intracellular ion concentrations. Prog Biophys Mol Biol. 2007;94(3):336–372. doi:10.1016/j.pbiomolbio.2006.10.001

59. Fraser JA, Middlebrook CE, Usher-Smith JA, Schwiening CJ, Huang CLH. The effect of intracellular acidification on the relationship between cell volume and membrane potential in amphibian skeletal muscle. Journal of Physiology. 2005;563(3):745–764. doi:10.1113/jphysiol.2004.079657

60. Eaton JW, Bateman D, Hauberg S. GNU Octave version 7.1.0 manual: a high-level interactive language for numerical computations. Published online 2022. https://www.gnu.org/software/octave/doc/v7.1.0/

61. Kay AR. How cells can control their size by pumping ions. Front Cell Dev Biol. 2017;5(MAY):1–14. doi:10.3389/fcell.2017.00041

62. Lew VL, Bookchin RM. Volume, pH and Ion-content regulation in human red cells. J Membrane Biol. 1986;92:57–74.

63. Barbosa NSV, Lima ERA, Boström M, Tavares FW. Membrane potential and ion partitioning in an erythrocyte using the Poisson-Boltzmann equation. Journal of Physical Chemistry B. 2015;119(21):6379–6388. doi:10.1021/acs.jpcb.5b02215

64. Jakobsson E. Interactions of cell volume, membrane potential, and membrane transport parameters. Am J Physiol Cell Physiol. 1980;7(3). doi:10.1152/ajpcell.1980.238.5.c196

65. Garrahan PJ, Rega AF. Cation loading of red blood cells. J Physiol. 1967;193(2):459–466. doi:10.1113/jphysiol.1967.sp008371

66. Gallagher PG. Disorders of erythrocyte hydration. Blood. 2017;130(25):2699–2708. doi:10.1182/blood-2017-04-590810

67. Dyrda A, Cytlak U, Ciuraszkiewicz A, et al. Local membrane deformations activate Ca2+-dependent K + and anionic currents in intact human red blood cells. PLoS One. 2010;5(2). doi:10.1371/journal.pone.0009447

68. Nardid OA, Schetinskey MI, Kucherenko Y V. Dimethyl sulfoxide at high concentrations inhibits non-selective cation channels in human erythrocytes. Gen Physiol Biophys. 2013;32(01):23–32. doi:10.4149/gpb_2013004

69. Desai SA, Bezrukov SM, Zimmerberg J. A voltage-dependent channel involved in nutrient uptake by red blood cells infected with the malaria parasite. Nature. 2000;406(6799):1001–1005. doi:10.1038/35023000

70. Richter S, Hamann J, Kummerow D, Bernhardt I. The monovalent cation “leak” transport in human erythrocytes: an electroneutral exchange process. Biophys J. 1997;73(2):733–745. doi:10.1016/S0006-3495(97)78106-2

71. Zade-Oppen AMM. The Effect of Mannitol, Sucrose, Raffinose and Dextran on Posthypertonic Hemolysis. Acta Physiol Scand. 1968;74(1-2):195–206. doi:10.1111/j.1365-201x.1968.tb10914.x

72. Miller RH, Mazur P. Survival of frozen-thawed human red cells as a function of cooling and warming velocities. Cryobiology. 1976;13(4):404–414. doi:10.1016/0011-2240(76)90096-1

73. Ogata N, Yoshii M, Narahashi T. Psychotropic drug block voltage-gated ion channels in neuroblastoma cells. Brain Res. 1989;476(1):140–144. doi:10.1016/0006-8993(89)91546-1

74. Nakazawa K, Ito K, Koizumi S, Ohno Y, Inoue K. Characterization of inhibition by haloperidol and chlorpromazine of a voltage-activated K+ current in rat phaeochromocytoma cells. Br J Pharmacol. 1995;116(6):2603–2610. doi:10.1111/j.1476-5381.1995.tb17214.x

75. Semionova YA, Zemlyanskikh NG, Orlova N V., Shpakova NM. Antihemolytic Efficiency of Chlorpromazine under Posthypertonic Shock and Glycerol Removal from Erythrocytes after Thawing. Problems of Cryobiology and Cryomedicine. 2017;27(1):051–060. doi:10.15407/cryo27.01.051

76. Rudenko S V., Patelaros S V. Cation-sensitive pore formation in rehydrated erythrocytes. BBA - Biomembranes. 1995;1235(1):1–9. doi:10.1016/0005-2736(94)00275-T

77. Patelaros S V. Influence of divalent cations Ca2+ and Zn2+ on the activation of posthypertonic hemolysis of human erythrocytes. Biopolym Cell. 1999;15(1):43–48. doi:10.7124/bc.000504

78. Gárdos G. The function of calcium in the potassium permeability of human erythrocytes. Biochim Biophys Acta. 1958;30(3):653–654. doi:10.1016/0006-3002(58)90124-0

79. Hoffman JF, Joiner W, Nehrke K, Potapova O, Foye K, Wickrema A. The hSK4 (KCNN4) isoform is the Ca2+-activated K+ channel (Gardos channel) in human red blood cells. Proc Natl Acad Sci U S A. 2003;100(12):7366–7371. doi:10.1073/pnas.1232342100

80. Föller M, Lang F. Ion Transport in Eryptosis, the Suicidal Death of Erythrocytes. Front Cell Dev Biol. 2020;8(July):1–9. doi:10.3389/fcell.2020.00597

81. Lang KS, Duranton C, Poehlmann H, et al. Cation channels trigger apoptotic death of erythrocytes. Cell Death Differ. 2003;10(2):249–256. doi:10.1038/sj.cdd.4401144

82. Amico-Ruvio SA, Murthy SE, Smith TP, Popescu GK. Zinc effects on NMDA receptor gating kinetics. Biophys J. 2011;100(8):1910–1918. doi:10.1016/j.bpj.2011.02.042

83. Mazur P, Rigopoulos N. Contributions of unfrozen fraction and of salt concentration to the survival of slowly frozen human erythrocytes: Influence of warming rate. Cryobiology. 1983;20(3):274–289. doi:10.1016/0011-2240(83)90016-0

84. Tenchini ML, Bolognani L, De Carli L. Effect of hypertonicity on survival of unprotected human cultured cells following freezing and thawing. Cryobiology. 1980;17(2):120–124. doi:10.1016/0011-2240(80)90015-2

85. Ishiguro H, Rubinsky B. Mechanical Interactions between Ice Crystals and Red Blood Cells during Directional Solidification. Cryobiology. 1994;31(5):483–500. doi:10.1006/cryo.1994.1059

86. Takamatsu H, Rubinsky B. Viability of deformed cells. Cryobiology. 1999;39(3):243–251. doi:10.1006/cryo.1999.2207

87. Fang XZ, Zhou T, Xu JQ, et al. Structure, kinetic properties and biological function of mechanosensitive Piezo channels. Cell Biosci. 2021;11(1):1–20. doi:10.1186/s13578-020-00522-z

88. Shpakova NM, Orlova N V., Nipot EY. Dehydration of mammalian erythrocytes affects their sensitivity to mechanical stress. Problems of Cryobiology and Cryomedicine. 2015;25(1):24–32. doi:10.15407/cryo25.01.024

89. Steponkus PL, Wiest SC. PLASMA MEMBRANE ALTERATIONS FOLLOWING COLD ACCLIMATION AND FREEZING. In: Plant Cold Hardiness and Freezing Stress. Elsevier; 1978:75–91. doi:10.1016/B978-0-12-447650-9.50011-3

90. Muldrew K, McGann LE. Mechanisms of intracellular ice formation. Biophys J. 1990;57(3):525–532. doi:10.1016/S0006-3495(90)82568-6

91. Muldrew K, McGann LE. The osmotic rupture hypothesis of intracellular freezing injury. Biophys J. 1994;66(2):532–541. doi:10.1016/S0006-3495(94)80806-9

92. Zheng S, Krump NA, McKenna MM, et al. Regulation of erythrocyte Na/K/2Cl cotransport by an oxygen-switched kinase cascade. Journal of Biological Chemistry. 2019;294(7):2519–2528. doi:10.1074/jbc.RA118.006393

93. Kahle KT, Rinehart J, Lifton RP. Phosphoregulation of the Na-K-2Cl and K-Cl cotransporters by the WNK kinases. Biochim Biophys Acta Mol Basis Dis. 2010;1802(12):1150–1158. doi:10.1016/j.bbadis.2010.07.009

94. Duhm J. Furosemide-sensitive K+ (Rb+) transport in human erythrocytes: Modes of operation, dependence on extracellular and intracellular Na+, kinetics, pH dependency and the effect of cell volume and N-ethylmaleimide. J Membr Biol. 1987;98(1):15–32. doi:10.1007/BF01871042

95. Adragna NC, Tosteson DC. Effect of volume changes on ouabain-insensitive net outward cation movements in human red cells. J Membr Biol. 1984;78(1):43–52. doi:10.1007/BF01872531

96. O’Neill WC, Mikkelsen RB. Furosemide-sensitive Na+ and K+ transport and human erythrocyte volume. Biochimica et Biophysica Acta (BBA) - Biomembranes. 1987;896(2):196–202. doi:10.1016/0005-2736(87)90180-5

97. Semplicini A, Spalvins A, Canessa M. Kinetics and stoichiometry of the human red cell Na+/H+ exchanger. J Membr Biol. 1989;107(3):219–228. doi:10.1007/BF01871937

98. Cahalan SM, Lukacs V, Ranade SS, Chien S, Bandell M, Patapoutian A. Piezo1 links mechanical forces to red blood cell volume. Elife. 2015;4(MAY):1–12. doi:10.7554/eLife.07370

99. Föller M, Kasinathan RS, Koka S, et al. TRPC6 contributes to the Ca 2+ leak of human erythrocytes. Cellular Physiology and Biochemistry. 2008;21(1-3):183–192. doi:10.1159/000113760

100. Belkacemi A, Fecher-Trost C, Tinschert R, et al. The TRPV2 channel mediates Ca2+ influx and the Δ9-THC-dependent decrease in osmotic fragility in red blood cells. Haematologica. 2021;106(8):2246–2250. doi:10.3324/haematol.2020.274951

101. Egée S, Kaestner L. The Transient Receptor Potential Vanilloid Type 2 (TRPV2) Channel–A New Druggable Ca2+ Pathway in Red Cells, Implications for Red Cell Ion Homeostasis. Front Physiol. 2021;12(June):1–5. doi:10.3389/fphys.2021.677573

102. Muraki K, Iwata Y, Katanosaka Y, et al. TRPV2 Is a Component of Osmotically Sensitive Cation Channels in Murine Aortic Myocytes. Circ Res. 2003;93(9):829–838. doi:10.1161/01.RES.0000097263.10220.0C

103. Makhro A, Hänggi P, Goede JS, et al. N-methyl-D-aspartate receptors in human erythroid precursor cells and in circulating red blood cells contribute to the intracellular calcium regulation. Am J Physiol Cell Physiol. 2013;305(11). doi:10.1152/ajpcell.00031.2013

104. Kaestner L, Bollensdorff C, Bernhardt I. Non-selective voltage-activated cation channel in the human red blood cell membrane. Biochim Biophys Acta Biomembr. 1999;1417(1):9–15. doi:10.1016/S0005-2736(98)00240-5

105. Bennekou P. The voltage-gated non-selective cation channel from human red cells is sensitive to acetylcholine. Biochimica et Biophysica Acta (BBA) - Biomembranes. 1993;1147(1):165–167. doi:10.1016/0005-2736(93)90328-W

106. Noël J, Zimmermann K, Busserolles J, et al. The mechano-activated K± channels TRAAK and TREK-1 control both warm and cold perception. EMBO Journal. 2009;28(9):1308–1318. doi:10.1038/emboj.2009.57

107. Stewart GW, Ellory JC, Klein RA. Increased human red cell cation passive permeability below 12 °C. Nature. 1980;286(5771):403–404. doi:10.1038/286403a0

108. Mironescu S. Hyperosmotic injury in mammalian cells. 3. Volume and alkali cation alterations of CHO cells in unprotected and DMSO-treated cultures. Cryobiology. 1978;15(2):178–191. doi:10.1016/0011-2240(78)90022-6

109. Semionova EA, Chabanenko EA, Orlova N V., Zubov PM, Shpakova NM. About mechanism of antihemolitic action of chlorpromazine under posthypertonic stress in erythrocytes. Problems of Cryobiology and Cryomedicine. 2017;27(3):219–229. doi:10.15407/cryo27.03.219

110. Kaestner L, Christophersen P, Bernhardt I, Bennekou P. The non-selective voltage-activated cation channel in the human red blood cell membrane: reconciliation between two conflicting reports and further characterisation. Bioelectrochemistry. 2000;52(2):117–125. doi:10.1016/S0302-4598(00)00110-0

111. Han B, Bischof JC. Direct cell injury associated with eutectic crystallization during freezing. Cryobiology. 2004;48(1):8–21. doi:10.1016/j.cryobiol.2003.11.002

112. Takamatsu H, Zawlodzka S. Contribution of extracellular ice formation and the solution effects to the freezing injury of PC-3 cells suspended in NaCl solutions. Cryobiology. 2006;53(1):1–11. doi:10.1016/j.cryobiol.2006.03.005

113. Klbik I, Čechová K, Milovská S, et al. Cryoprotective Mechanism of DMSO Induced by the Inhibitory Effect on Eutectic NaCl Crystallization. J Phys Chem Lett. 2022;13(48):11153–11159. doi:10.1021/acs.jpclett.2c03003

114. Huang H, Zhao G, Zhang Y, Xu J, Toth TL, He X. Predehydration and Ice Seeding in the Presence of Trehalose Enable Cell Cryopreservation. ACS Biomater Sci Eng. 2017;3(8):1758–1768. doi:10.1021/acsbiomaterials.7b00201

115. Shen L, Guo X, Ouyang X, Huang Y, Gao D, Zhao G. Fine-tuned dehydration by trehalose enables the cryopreservation of RBCs with unusually low concentrations of glycerol. J Mater Chem B. 2021;9(2):295–306. doi:10.1039/d0tb02426k

116. Klbik I, Čechová K, Milovská S, et al. Polyethylene glycol 400 enables plunge-freezing cryopreservation of human keratinocytes. J Mol Liq. 2023;379:121711. doi:10.1016/j.molliq.2023.121711

117. Knorpp CT, Merchant WR, Gikas PW, Spencer HH, Thompson NW. Hydroxyethyl starch: Extracellular cryophylactic agent for erythrocytes. Science (1979). 1967;157(3794):1312–1313. doi:10.1126/science.157.3794.1312

118. Horn EP, Sputtek A, Standl T, Rudolf B, Kühnl P, Schulte Am Esch J. Transfusion of autologous, hydroxyethyl starch-cryopreserved red blood cells. Anesth Analg. 1997;85(4):739–745. doi:10.1097/00000539-199710000-00006

119. Spindler R, Rosenhahn B, Hofmann N, Glasmacher B. Video analysis of osmotic cell response during cryopreservation. Cryobiology. 2012;64(3):250–260. doi:10.1016/j.cryobiol.2012.02.008

120. Kellenberger S, Schild L. Epithelial sodium channel/degenerin family of ion channels: A variety of functions for a shared structure. Physiol Rev. 2002;82(3):735–767. doi:10.1152/physrev.00007.2002

121. Stachecki JJ, Cohen J, Willadsen SM. Cryopreservation of Unfertilized Mouse Oocytes: The Effect of Replacing Sodium with Choline in the Freezing Medium. Cryobiology. 1998;37(4):346–354. doi:10.1006/cryo.1998.2130

122. Stachecki JJ, Willadsen SM. Cryopreservation of Mouse Oocytes Using a Medium with Low Sodium Content: Effect of Plunge Temperature. Cryobiology. 2000;40(1):4–12. doi:10.1006/cryo.1999.2215

123. Quintans CJ. Birth of two babies using oocytes that were cryopreserved in a choline-based freezing medium. Human Reproduction. 2002;17(12):3149–3152. doi:10.1093/humrep/17.12.3149

124. Talevi R, Barbato V, Mollo V, et al. Replacement of sodium with choline in slow-cooling media improves human ovarian tissue cryopreservation. Reprod Biomed Online. 2013;27(4):381–389. doi:10.1016/j.rbmo.2013.07.003

125. Lawonn P, Hoffmann EK, Hougaard C, Wehner F. A cell shrinkage-induced non-selective cation conductance with a novel pharmacology in Ehrlich-Lettre-ascites tumour cells. FEBS Lett. 2003;539(1-3):115–119. doi:10.1016/S0014-5793(03)00210-2

126. Takahashi T, Williams RJ. Thermal shock hemolysis in human red cells. Cryobiology. 1983;20(5):507–520. doi:10.1016/0011-2240(83)90039-1

127. Woolgar AE, Morris GJ. Some combined effects of hypertonic solutions and changes in temperature on posthypertonic hemolysis of human red blood cells. Cryobiology. 1973;10(1):82–86. doi:10.1016/0011-2240(73)90011-4

